# The YdiU Domain Modulates Bacterial Stress Signaling through Mn^2+^-dependent UMPylation

**DOI:** 10.1101/692707

**Authors:** Yinlong Yang, Yingying Yue, Nannan Song, Cuiling Li, Zenglin Yuan, Yan Wang, Yue Ma, Hui Li, Fengyu Zhang, Weiwei Wang, Haihong Jia, Peng Li, Xiaobing Li, Qi Wang, Zhe Ding, Hongjie Dong, Lichuan Gu, Bingqing Li

## Abstract

Sensing stressful conditions and adjusting cellular metabolism to adapt to the environment is essential for bacteria to survive in variable situations. Here, we describe a new stress-related protein YdiU, and characterize YdiU as an enzyme that catalyzes the covalent attachment of uridine 5’-monophosphate to a protein tyrosine/histidine residue—a novel modification defined as UMPylation. Mn^2+^ serves as an essential co-factor for YdiU-mediated UMPylation. UTP and Mn^2+^-binding converts YdiU to an aggregate-prone state facilitating the recruitment of chaperones. The UMPylation of chaperones prevents them from binding co-factors or clients, thereby impairing their function. Consistent with the recent finding that YdiU acts as an AMPylator, we further demonstrate that the self-AMPylation of YdiU padlocks its chaperone-UMPylation activity. The detailed mechanism is proposed based on Apo-YdiU, YdiU-ATP, YdiU-AMP crystal structures and molecular dynamics simulation models of YdiU-UTP and YdiU-UTP-peptide. *In vivo* data demonstrate that YdiU effectively protects *Salmonella* from stress-induced ATP depletion through UMPylation.

**Highlights:**

1. YdiU involves in stress-resistance of *Salmonella*.
2. YdiU mediates protein UMPylation in a Mn^2+^-dependent manner.
3. Structural insights into YdiU-mediated UMPylation.
4. UMPylation of chaperones by YdiU modulates their function.

## INTRODUCTION

Bacteria survive in a changeable environment and have evolved myriad mechanisms to sense and respond to stress conditions. The most well-known strategy is to change gene expression in response to environmental variation(Browning and Busby, 2004). Transcription factors or alternative sigma factors are activated under stress conditions, and bind to their target genes to activate or inhibit transcription of the regulated genes(Balleza et al., 2009; Helmann and Chamberlin, 1988). Post-translational modifications (PTMs) of proteins also play an important role in the regulation of stress response. Bacterial two-component signal transduction systems, consist of a histidine kinase and a response regulator, regulate cellular physiology through a series of phosphorylation events(Hoch, 2000; Stock et al., 2000). Protein acetylation modulates many cellular processes of bacteria such as metabolism, bacterial chemotaxis and DNA replication response to different environments(Carabetta and Cristea, 2017; Hu et al., 2010; Jones and O’Connor, 2011). Additionally, other modifications such as carbonylation, nitration, N-myristoylation, phospholipid, glycosylation, ADP-ribosylation, and AMPylation have been detected in bacteria, however, the targets and mechanism of those modifications and their role in stress resistance remain largely unknown(Cain et al., 2014; Dukan and Nystrom, 1998; Grangeasse et al., 2015; Schwarz and Aebi, 2011).

The YdiU (also known as SelO or UPF0061) domain is a pseudokinase domain that is widely found in the three kingdoms of life and is highly conserved from *E.coli* to human (43% full-length sequence identity)(Dudkiewicz et al., 2012). Over 15,000 proteins contain a predicted YdiU domain (InterPro Database; IPR003846; http://www.ebi.ac.uk/interpro/index.html). However, the *in vivo* and *in vitro* function of the YdiU domain has not been characterized until recently, making it one of “the top ten most-attractive unknown domains”(Galperin and Koonin, 2004, 2010). Most species of bacteria contain a single YdiU domain protein, and previous transcriptomic data showed an increase in YdiU expression during the recovery stage after heat shock, hydrostatic pressure, and osmotic stress implying a relationship between YdiU and bacterial stress resistance(Erasmus et al., 2003; Hsu-Ming et al., 2012; Phadtare and Inouye, 2004). Most recently, Sreelatha *etal* reported a role of YdiU in redox homeostasis and identified YdiU as an AMPylator that can transfer AMP from ATP to Ser, Thr, and Tyr residues on protein substrates(Sreelatha et al., 2018). In addition to auto-AMPylation of YdiU, two proteins involved in redox homeostasis were also found to be AMPylated substrates of YdiU. The discovery by Sreelatha *et al*. that YdiU acts as an enzyme that mediates protein modification significantly advances investigation into the functions of YdiU family proteins. However, the details of how AMPylation regulates the activities of substrates and whether YdiU is able to modify other protein substrates and functions as part of other responses requires further investigation.

In this study, we demonstrated a role of YdiU in bacterial stress resistance and discovered an unexpected activity of YdiU to catalyze the UMP-modification of bacterial chaperones. Chaperones play an important role under stress conditions by repairing misfolded proteins. In bacterial chaperone networks, GroEL and DnaK function as central hubs, with contributions of activity from other proteins, such as HtpG and ClpB(Gragerov et al., 1992; Thomas and Baneyx, 2000). Our data clearly revealed that GroEL, DnaK, HtpG, and ClpB, could all be UMPylated by YdiU both *in vivo* and *in vitro*. Interestingly, YdiU is a high-affinity manganese-binding protein and the UMPylation activity of YdiU depends on Mn^2+^. In the presence of Mn^2+^, the affinity of YdiU to bind UTP is one hundred times greater than that for ATP. More importantly, UTP and Mn^2+^ binding results in the exposure of the hydrophobic region of YdiU that is essential for chaperone recruitment. The detailed mechanism of YdiU-mediated NMPylation is proposed by determining the crystal structure of Apo-YdiU, YdiU-ATP-Mg^2+^, YdiU-ATP-Mn^2+^, and YdiU-AMP-Mg^2+^ complexes and a structural model of YdiU-UTP-Mn^2+^ obtained by molecular dynamics simulation. Further, we revealed that the UMPylation of chaperones by YdiU prevented the binding of the modified chaperones to either downstream co-factors or their substrates. Chaperone UMPylation was inhibited by self-AMPylation that senses the intracellular level of ATP. *In vivo* data show that *Salmonella* benefits from YdiU-mediated UMPylation under severe heat injury and UMPylation occurs in response to heat stress in a YdiU-dependent manner.

## RESULTS

### The Expression of YdiU is Stress-dependent in *Salmonella*

To investigate the conditions under which *Salmonella* YdiU is expressed, the mRNA and protein levels of YdiU were measured in *Salmonella* cultured under different conditions by q-PCR and western blot. Interestingly, YdiU was barely expressed when *Salmonella* was cultured in Luria-Bertani medium at 37°C (normal laboratory growth condition), but increased to high level under multiple stress conditions such as nutritional deficiency, H_2_O_2_ stress, and acid stress (Fig.1A, B). Then the expression of YdiU were detected after *Salmonella* culture was incubated at 42-55°C for 5 min. Compared with the cells cultured in 37°C, the level of YdiU was observably induced by heat shock and accumulated with increased heat time (Fig.1C-F). The finding that stress conditions significantly elevate expression of YdiU suggests that YdiU may act in the survival of *Salmonella* under stress conditions.

**Fig. 1.**
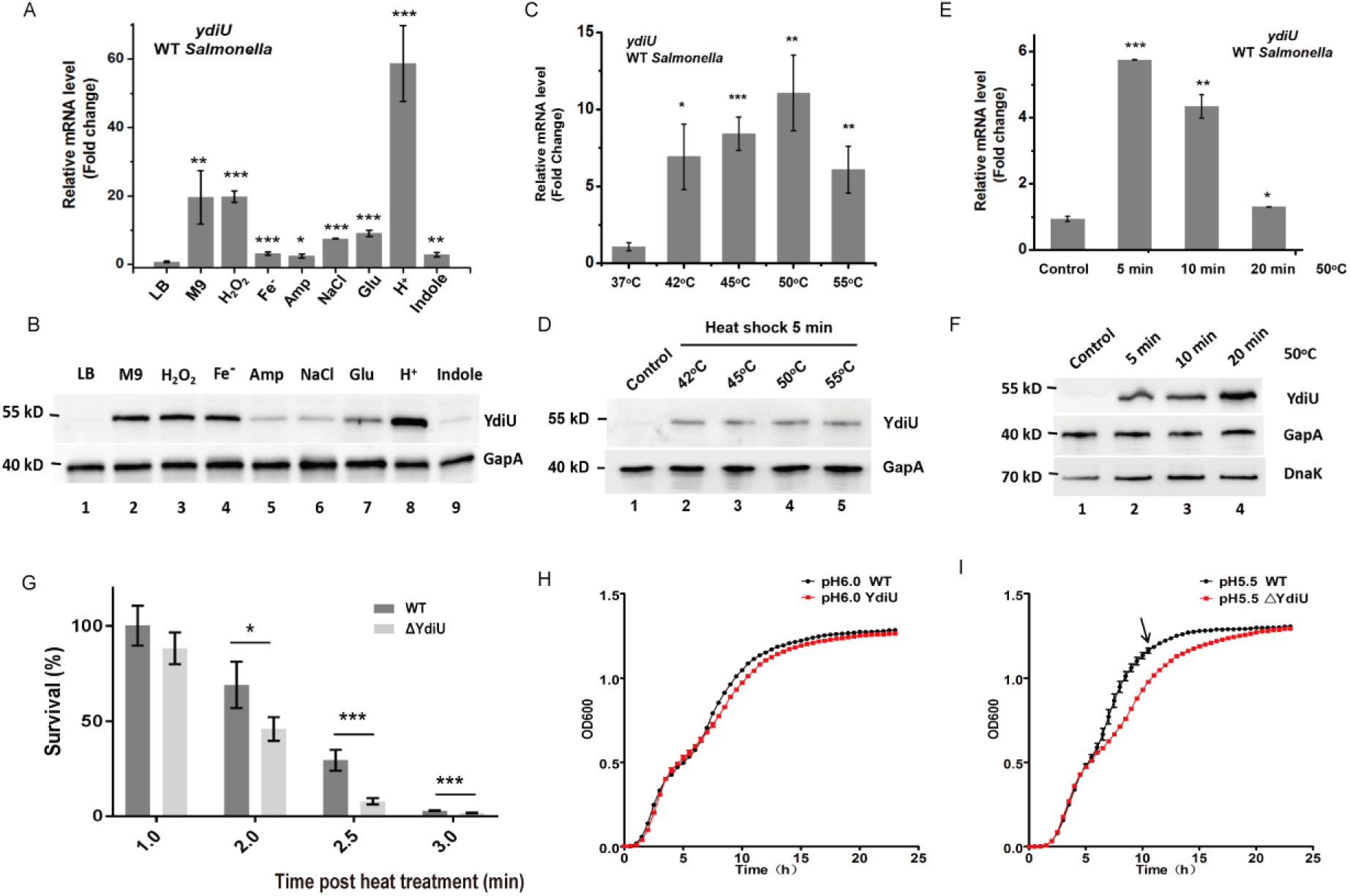
YdiU is involved in stress-resistance of *Salmonella.* (A-B) The transcription and protein levels of YdiU in *Salmonella* cultivated under different conditions were detected by qRT-PCR and western blot respectively. GapA (also known as GADPH) was used as a loading control. (C-D) The transcription and protein level of YdiU in *Salmonella* 30 min post-heat treatment at the indicated temperature for 5min. The cells cultured at 37°C were used as a control. (E-F) The transcription and protein level of YdiU in *Salmonella* 30 min post-heat treatment after 5-20 min heat treatment at 50°C. (G) Survival ratios of WT and ΔYdiU were detected following treatment at 55°C for the indicated time. (H-I) The growth curves of WT and ΔYdiU cultivated in LB medium with different pH. The above experiments were performed as three or four replicates and the mean values are presented. The statistical significance is indicated by ***P < 0.001, **P < 0.005, *P < 0.05, as compared to control.

### YdiU Protects *Salmonella* from Multiple Stressful Conditions

To explore the function of YdiU, we constructed a YdiU knockout strain of *Salmonella* (ΔYdiU). The tolerance of stressful conditions was tested in the wild-type *Salmonella* and the ΔYdiU strain (Fig.1G-L). Compared with the WT strain, the ΔYdiU strain exhibited significantly reduced survival rate after a long heat treatment, suggesting that YdiU protects *Salmonella* against heat-induced cell death, especially caused by severe heat stress (Fig 1G). An apparent growth defect of the ΔYdiU strain is observed in the late logarithmic phase when cultivated in low pH medium, which is not observed for wild type *Salmonella* (Fig.1H-I), further confirming the important role of YdiU in stress tolerance.

### UMPylated Proteins Identified in YdiU-expressing *Salmonella*

To explore the function of YdiU, an expression plasmid of YdiU was transformed into the ΔYdiU strain to generate a strain that expressed YdiU under tight regulation (pYdiU). The production of YdiU in the pYdiU strain was determined for different concentrations of inducer L-arabinose (Supplementary Fig.S1). The ΔYdiU and pYdiU strains were cultured in LB medium with 0.1% L-arabinose and used to study the phenotype of *Salmonella* in the presence or absence of YdiU. Mass spectrometry-based proteomic analysis was performed for the ΔYdiU and pYdiU strains. Only a few proteins showed a more than 50% change in expression suggesting that YdiU might not act widely as a regulator at the level of protein expression. Recently, YdiU was identified as an AMPylator that can modify substrates with adenine 5’-monophosphate (AMP) in *E.coli*, so we analyzed the protein modifications in ΔYdiU and pYdiU *Salmolnella* using ProteinPilot software^19^. Surprisingly, no AMPylated peptides were detected in YdiU-expressing *Salmonella*, however, fifty-six peptides that were covalently modified with an uridine 5’-monophosphate (UMP) molecule were detected in YdiU-expressing *Salmonella* (Supplementary Table S1). Previously, UMPylation was only observed in glutamine synthetase system, in which the synthesis rate of glutamine is modulated by self-modification between subunits(Adler et al., 1975). Because the YdiU domain includes a nucleotide binding site, we hypothesized that YdiU could act as an UMPylator (an enzyme that catalyzes UMPylation) *in vivo* (Fig.2E). UMPylated peptides identified in YdiU-expressing *Salmonella* were mapped to forty-six different *Salmonella* proteins involved in multiple life activities (Fig.2A, B and Supplementary Table S2). Analysis of the primary sequences of the identified UMPylated peptides reveals obvious common characteristics (Fig.2C, D): i). UMPylation was observed only on a tyrosine or histidine residue; ii). Most UMPylation sites (41/56) are close to one or more negatively charged residues (Asp or Glu); and iii). There was enrichment of hydrophobic residues around the UMPylation sites.

**Fig. 2.**
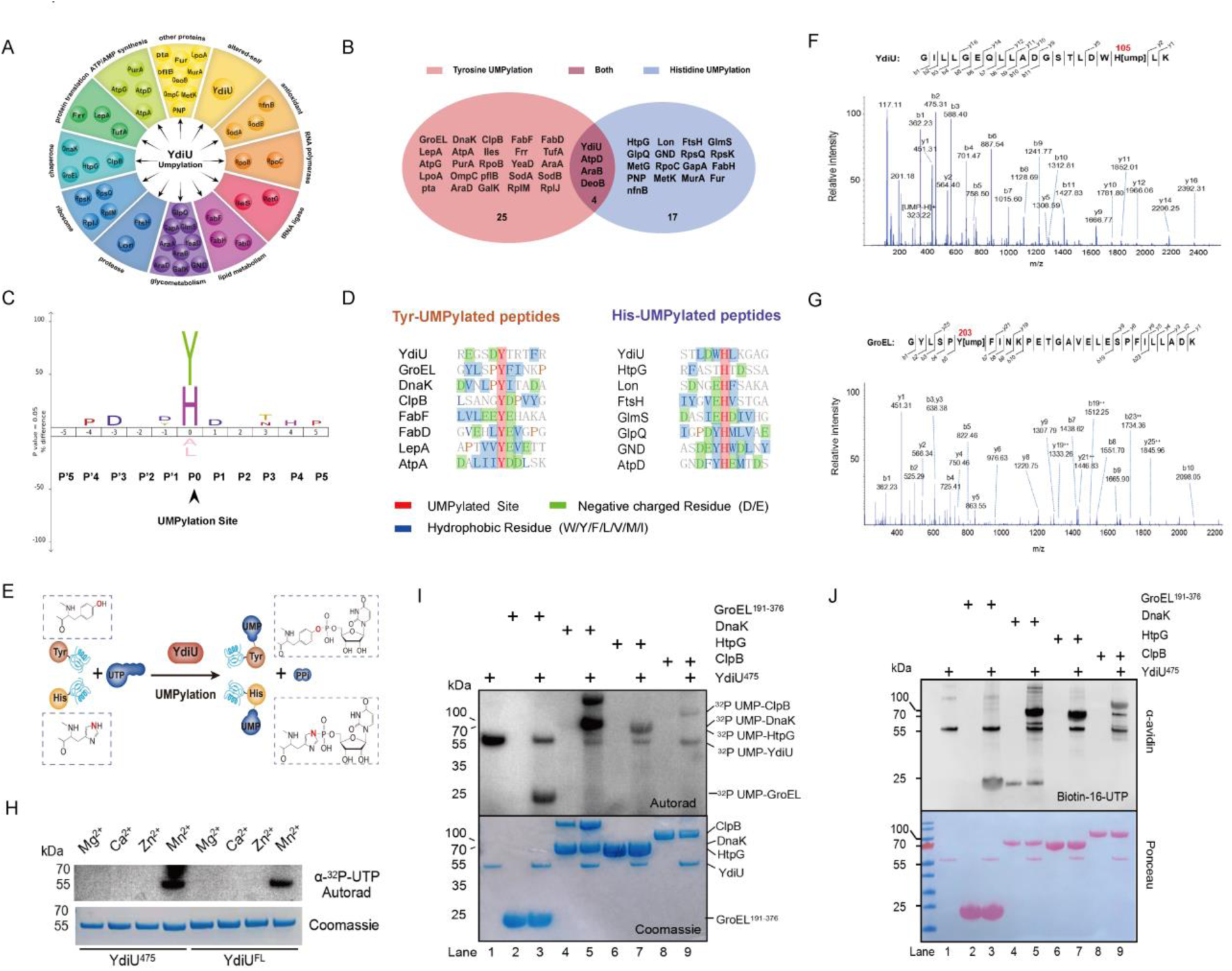
YdiU mediates UMPylation *in vivo* and *in vitro.* (A) Schematic of UMPylated proteins identified in YdiU-expressing *Salmonella* classified according to their function. (B) Venn diagram of the identified UMPylated proteins grouped by Tyrosine or Histidine UMPylation. (C) Sequence profiles for UMPylated peptides aligned with the iceLogo server. Significant conserved sequence patterns are highlighted. The *Salmonella* genome was set as the background (reference) sequence. (D) The sequence signature of eight Tyr-UMPylated peptides and eight His-UMPylated peptides. UMPylated residues are shown in red, the negative charged residues and hydrophobic residues, are highlighted in green and blue, respectively. (E) Diagram for YdiU-mediated UMPylation. (F-G) Electrospray ionization MS/MS spectra of UMPylated peptides identified in vivo. The b and y ions are marked and indicated along the peptide sequence above the spectra. (F) Histidine-UMPylated peptide from YdiU. (G) Tyrosine-UMPylated peptide from GroEL. (H) *In vitro* assays of self-UMPylation of YdiU with different metal ions. YdiU^fl^ and YdiU^475^ were incubated with α^32^P-UTP and corresponding divalent ions, followed by SDS-PAGE and autoradiography. (I) *In vitro* UMPylation of chaperones mediated by YdiU using α^32^P-UTP as donor. UMPylation assays were performed with YdiU^475^ and the indicated recombinant candidate substrates in a reaction buffer containing α^32^P-UTP and MnCl_2_. (J) *In vitro* UMPylation of chaperones mediated by YdiU using biotin-16-UTP as donor. UMPylation assays were performed with YdiU^475^ and the indicated recombinant candidate substrates with biotin-16-UTP and MnCl_2_ followed by avidin blotting. Total proteins were visualized by Ponceau S staining of the blot before blocking.

The UMPylated proteins in YdiU-expressing *Salmonella* included YdiU, suggesting that this protein can catalyze self-UMPylation (Fig.2F and Supplementary Fig.S2A). Other substrates of YdiU are key players involved in many vital activities, revealing a potentially extensive regulatory role of YdiU and UMPylation (Fig.2A). Remarkably, all major bacterial chaperones (GroEL, DnaK, HtpG, and ClpB) were UMPylated by YdiU *in vivo* (Fig.2G and Supplementary Fig.S2B-D). Chaperones are essential for bacteria to survive under stress conditions due to their activity to repair misfolded proteins, so the regulation of chaperones by YdiU through UMPylation may be associated with stress resistance of *Salmonella* when subjected to stressful conditions. Thus, chaperones were focused on and examined in more detailed in this study.

### YdiU Catalyzes *in vitro* UMPylation in a Mn^2+^-dependent Manner

To further confirm the function of YdiU as an UMPylator, we purified full-length *E.coli* YdiU (478aa, YdiU^fl^) and a truncated version lacking the C-terminal three amino acids (YdiU^475^). The purified proteins were tested using *in vitro* assays with ɑ^32^P-UTP in the presence of different divalent metal ions (Fig.2H). Interestingly, ^32^P-UMP-YdiU products were observed in reaction assays prepared with buffers containing Mn^2+^. *Salmonella* increases the absorption of manganese in response to multiple stress signals(Jakubovics and Jenkinson, 2001; Kehres et al., 2000), and the stimulation of UMPylation activity of YdiU by manganese suggests a regulatory mechanism of enzyme activity that utilizes metal ions. Subsequent *in vitro* UMPylation assays of chaperones were performed using Mn^2+^ as an activator and either ɑ^32^P-UTP or biotin-16-UTP as the UMP donor (Fig.2I, J). The assay results clearly showed that the tested chaperones could be UMPylated by YdiU *in vitro*.

### YdiU is A High-Affinity Manganese-Binding Protein

Our data showed that YdiU catalyzed UMPylation only in the presence of Mn^2+^, so we hypothesized that YdiU might specifically interact with Mn^2+^. To test this, we used microscale thermophoresis (MST) and surface plasmon resonance (SPR) to study the manganese-binding affinity of YdiU. Both MST and SPR experiments showed a strong affinity of YdiU and Mn^2+^, with calculated dissociation constants of 7∼10 μM (Fig.3A, B). The binding of YdiU and Mg^2+^ was also tested, however, no binding signal was observed (Fig.3A).

**Fig. 3.**
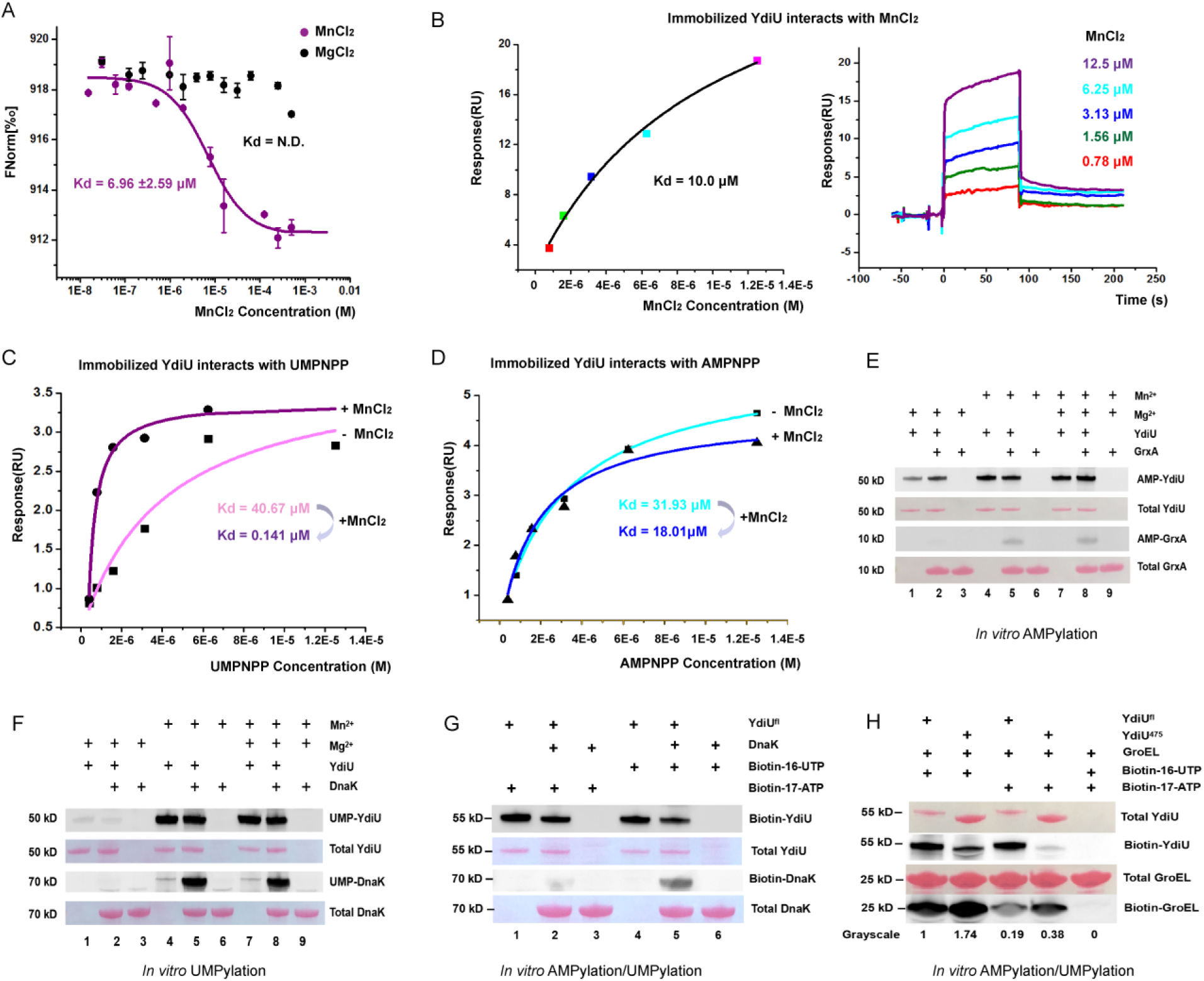
Mn^2+^ is essential for UTP preference and UMPyaltion activity. (A) Analysis of Mn^2+^ or Mg^2+^ binding affinity to YdiU using MST assay. (B) Analysis of Mn^2+^ or Mg^2+^ binding affinity to YdiU using SPR assay. (C-D) SPR analysis of the effects of Mn^2+^ on the affinity between YdiU and UTP or ATP analog. For the Mn^2+^-adding groups, 50 μM MnCl_2_ was added to the running buffer. (E-H) *In vitro* AMPylation/UMPyaltion assays using biotin-16-UTP or biotin-17-ATP as NMP donor. Biotin-labeled proteins were detected by avidin blotting and total proteins were visualized by Ponceau S staining. (E) *In vitro* AMPylation assay performed with GrxA and YdiU in the presence of 10 mM Mg^2+^ or Mn^2+^ or both. (F) *In vitro* UMPylation assay performed with DnaK and YdiU in presence of 10 mM Mg^2+^ or Mn^2+^ or both. (G) Comparison of AMPylation and UMPylation catalyzed by YdiU with DnaK in the presence of 10 mM Mn^2+^. (H) Comparison of AMPylation and UMPylation catalyzed by YdiU with GroEL^191-376^ in the presence of 10 mM Mn^2+^.

### Manganese Favors the Binding of YdiU to UTP

YdiU was first reported as an AMPylator, however, our *in vivo* characterization of YdiU-expressing *Salmonella* failed to identify substrates with AMPylation, though dozens of targets with UMPylation were detected. To examine the affinities of YdiU with its nucleotide-substrates, a series of SPR experiments were employed with non-degradable analogues of UTP (UMPNPP) or ATP (AMPNPP) (Fig.3C-D). We examined the interaction in the absence of Mn^2+^ and in presence of Mn^2+^. The obtained dissociation constant (K_d_) values for the interaction of YdiU and UMPNPP was ∼41 μM and that of YdiU and AMPNPP was ∼32 μM in the buffer without Mn^2+^. In the presence of 50 μM Mn^2+^, however, the Kd value for YdiU and UMPNPP interaction was decreased to ∼0.14 μM and that of YdiU-AMPNPP remained around 18 μM suggesting that Mn^2+^-binding favors UTP-binding of YdiU.

### Manganese is Essential for Substrate-UMPylation/AMPylation of YdiU

Sreelatha *etal* identified *E.coli* GrxA as the substrate of YdiU-mediated AMPylation. We purified *E.coli* GrxA and performed AMPylation experiments *in vitro*. Our data demonstrated that GrxA could be AMPylated by YdiU only if Mn^2+^ was present in the reaction system (lanes 5 and 8 in Fig.3E). Interestingly, the self-AMPylation of YdiU was not Mn^2+^-dependent (lanes 1 and 2 in Fig.3E). Consistent with previous experiments (Fig.2H-J), both self- and substrate-UMPylation mediated by YdiU were Mn^2+^-dependent processes (Fig.3F). To determine if the UMPylated substrates we identified *in vivo* could be AMPylated by YdiU *in vitro*, the AMPylation activity of YdiU was assayed in the presence of DnaK and GroEL (Fig.3G-H). The result showed that DnaK could not be AMPylated by YdiU *in vitro* even in presence of Mn^2+^. However, GroEL was able to be AMPylated by YdiU *in vitro,* but to a much lower extent than the observed UMPylation. The above data reveal the important role of Mn^2+^ in regulating the enzyme activity of YdiU and the striking preference for UMPylation by YdiU.

### UTP and Mn^2+^-binding Promotes an Aggregation-prone State for YdiU

To further investigate the mechanism of YdiU-associated UMPylation, we tried to obtain crystal structures of YdiU. Apo-YdiU crystalized successfully, however, YdiU complexed with UTP analog and Mn^2+^ precipitated in almost all the drops in the crystal trials (Supplementary Fig.S3A). We next tried to soak UTP analog and Mn^2+^ into crystals of Apo-YdiU, but those crystals became withered in the buffer containing UTP and Mn^2+^. We found that YdiU started to precipitate in the presence of a 10-fold excess of UTP, and this effect was aggravated when 5 mM MnCl_2_ was added (Supplementary Fig.S3B). The behavior of YdiU in the presence of UTP/ATP and Mn^2+^ was evaluated with a biologics stability screening platform (Uncle stability platform). DLS results showed two peaks exist in hydrodynamic diameter distributions in solutions of YdiU. The first one with a hydrodynamic diameter of ∼5.9 nm might be the YdiU monomer while the other one of ∼45 might be the aggregation of YdiU molecules. When incubated with UTP and Mn^2+^, YdiU tends to the aggregation state while exists in monomer form when in presence of ATP and Mn^2+^. (Supplementary Fig.S3C). The conformational stability was also confirmed by DSF, the Tm1 values of apo-YdiU, YdiU-UTP-Mn^2+^ and YdiU-ATP-Mn^2+^ are 38.3, 40.6 and 50.3 °C, respectively. (Supplementary Fig.S3D). All these results indicate that YdiU tends to an aggregation-prone state when bound to UTP and Mn^2+^ while tends to be stable monomer when bound to ATP and Mn^2+^.

### Structural Insights into the Catalytic Mechanism of YdiU-mediated NMPylation

Although we were unable to determine the crystal structure of YdiU-UTP-Mn, we successfully solved structures of Apo-YdiU, YdiU-ATP-Mn, and YdiU-ATP-Mg. Interestingly, a transient state of YdiU-AMP-PPi-Mg was also obtained. The overall structure of YdiU exhibits a conformation in which the catalysis domain is stabilized by the C-terminal regulatory domain, which looks like a big “clamp” (Fig.4A, B). The AMPPNP or AMP/PNP molecule is clearly defined in the corresponding electron density maps (Fig.4C, D). The nucleic acid binding site is located in a cleft at the interface between the N-lobe and C-lobe of the catalysis domain. The ATP analog is positioned with Mg^2+^ or Mn^2+^ in the active site via a novel binding mode in which the ɤ-phosphate is deeply buried in a pocket and the purine ring is located outside the pocket (Fig.4F-G, Supplementary Fig.S4A). The locations of Mg^2+^ and Mn^2+^ were identical in the obtained structures (Supplementary Fig.S4A). A structure of *P. syringae* YdiU in complex with AMPPNP, Mg^2+^ and Ca^2+^ was reported recently by Sreelatha *etal*^19^. Despite the sequence identity of 48%, the catalytic domains of *E.coli* YdiU and *P. syringae* YdiU displays a very similar conformation(Sreelatha et al., 2018). The basic group of AMPPNP is fixed through two hydrogen-bond interactions with Asp119 and Gly120 (Fig.4F). An extensive network of interactions orients the β and ɤ phosphates of AMPPNP with Arg87, Lys107, Arg170 and Arg177. Two Mg^2+^ coordinate with phosphate groups, and Asp 256 and Asn 247 clamp the AMPPNP to facilitate ATP hydrolysis (Fig.4G). Before binding to nucleotides, Asp256 interacts with Arg170 and Arg177 in the active center of apo-YdiU, and Arg87 interacts with Asp119 via electrostatic interaction (Fig.4E, Supplementary Fig.S4B). Superposition of the YdiU-AMPPNP and YdiU-AMP structures reveals that the purine ring and nucleic sugar adopt the same position in the active center, however, the phosphate of AMP is displaced from the α-phosphate of ATP and instead is coordinated with Gln75, Asn244, and Asn247 in the YdiU-AMP structure (Supplementary Fig.S4C and Fig.4G). Asp246 is the only basic residue near the α-phosphate of ATP, and we hypothesized that this residue might act as the general base for nucleophilic attack.

**Fig. 4.**
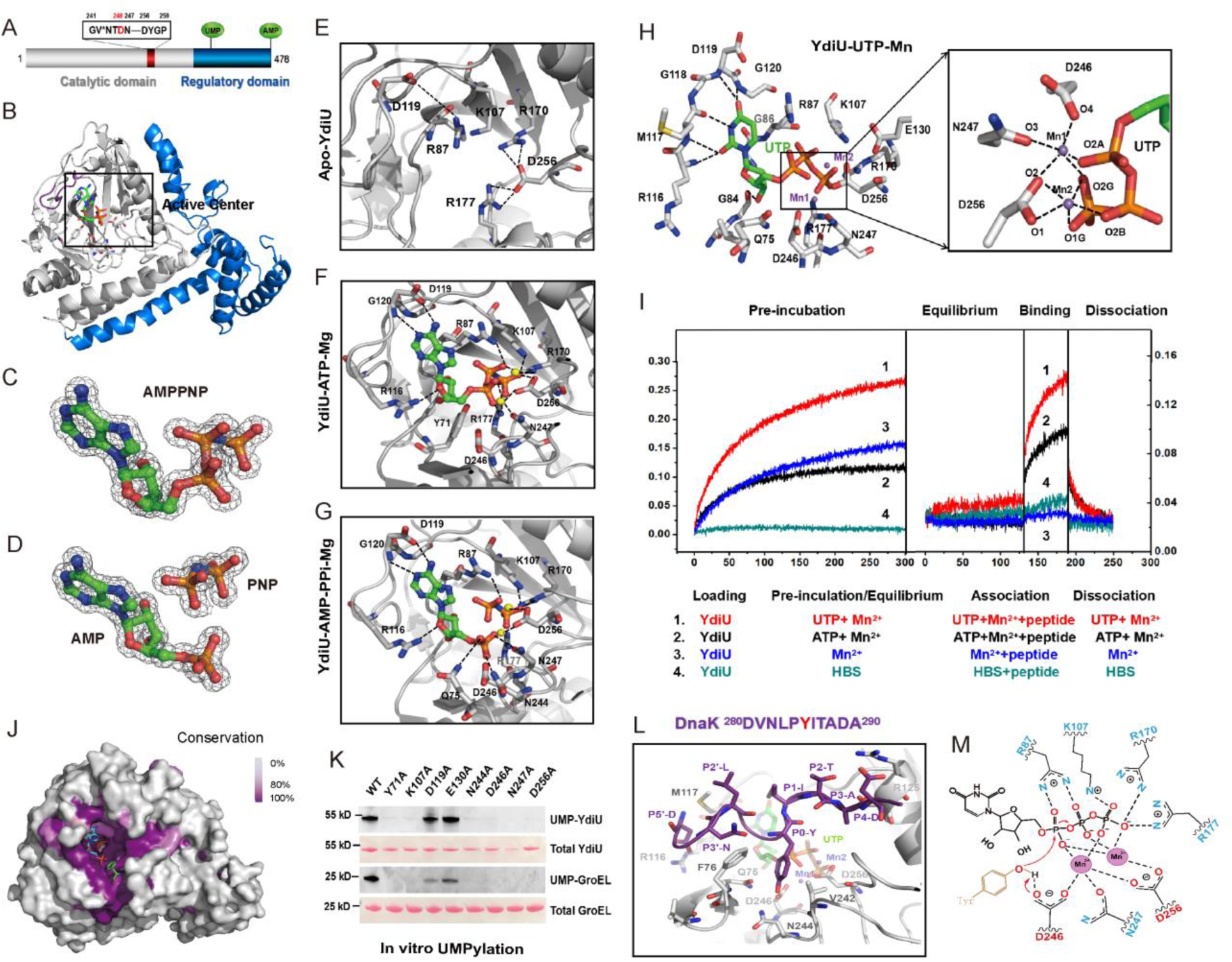
The catalytic mechanism of YdiU-mediated UMPylation. (A) Schematic representation of the domain architecture of YdiU. The conserved catalysis motif and the major self-UMPylation/self-AMPylation sites are highlighted. (B) The overall structure of YdiU-AMPPNP complex is shown in cartoon. The loop from Lys107 to Arg121 responsible for basic fixing is named the BF loop and is highlighted by purple. The active center is marked by a black frame. (C-D) Well-defined electron density maps of AMPPNP and AMP-PPi from YdiU-AMPNPP and YdiU-AMP-PPi, respectively. The 2F_o_-F_c_ omit electron densities are contoured at 2.0 σ. (E-G) Zoomed view of the active sites of Apo-YdiU, YdiU-AMPNPP and YdiU-AMP-PPi. Overall structure of YdiU is shown in cartoon and the key residues are highlighted in sticks. Hydrogen bonding interactions between residues of YdiU or YdiU and nucleotide are all indicated as black dashed lines. (H) Zoomed view of the active sites of the YdiU-UTP-Mn^2+^ structure generated by molecular dynamics simulation. (I) BLI analysis of the binding of YdiU to DnaK peptide DVNLPYITADA in the presence or absence of nucleotide and Mn^2+^. (J) Sequence conservation of YdiU among species. Overall structure of YdiU is shown in surface mode. Highly conserved residues are colored in purple and less conserved ones are colored in light grey. Substrate Tyr is docked into the predicted substrate-binding site and shown in stick form. (K) UMPylation activity of YdiU mutants with Biotin-16-UTP, as detected by streptavidin HRP blot. Total proteins were visualized by Ponceau S staining. (L) Model of YdiU-UTP-Mn-DnaK peptide generated by molecular docking. Also see Supplementary Fig.S6 and Fig.S7. (M) The proposed mechanism of YdiU-mediated UMPylation. Asp 246 acts as the general base and activates the oxygen of the hydroxyl group from Tyr for nucleophilic attack. Also see Supplementary Fig.S8.

### Structural Basis for the UTP Preference of YdiU

YdiU exhibits a preference for UTP in the presence of Mn^2+^ (Fig.3C,D). To investigate the mechanisms underlying the nucleotide selectivity of YdiU, we modeled the YdiU-UTP-Mn^2+^ and YdiU-UTP-Mg^2+^ complexes using molecular docking based on the crystal structure of Apo-YdiU. The docking result positioned UTP in the same cavity of YdiU as ATP. Next, structure-based molecular dynamics simulation was performed of YdiU binding with either UTP or ATP with Mn^2+^. The obtained convergence parameters suggested that both structures of YdiU-ATP and YdiU-UTP were stable during simulation. Six hydrogen bonds occurred between YdiU and ATP while eleven occurred in YdiU-UTP. The hydrogen-bond interactions in YdiU-UTP-Mn^2+^ and YdiU-ATP-Mn^2+^ systems are listed in Supplementary Table S3. The active center in YdiU-UTP-Mn^2+^ exhibits a very stable conformation that is stabilized by an interaction network between YdiU, UTP and Mn^2+^ (Fig.4H). The loop Gly110-Arg121 of YdiU showed a tight interaction with the pyrimidine ring of UTP through four hydrogen bonds (Fig.4H, Supplementary Table S3). Moreover, the coordination of Mn^2+^ with YdiU and the UTP molecule in the YdiU-UTP-Mn^2+^ model is more compact than that in the YdiU-ATP-Mn^2+^ model, and Asp246 provides an extra bond interaction with Mn^2+^ in the YdiU-UTP-Mn^2+^ model which is absent in the YdiU-ATP-Mn^2+^ model (Supplementary Fig.S4D-G). Sequence alignment shows that the active site residues are strictly conserved from bacteria to human, suggesting a uniform mechanism for YdiU homologues (Fig.4J, Supplementary Fig.S5). The exact role of these residues in UMPylation was investigated further by mutagenesis. Most mutations of residues in the active center completely abolished enzyme activities (Fig.4K). The above structural information well explains the phenomenon that the affinity of YdiU and UTP is dramatically improved by Mn^2+^. In conclusion, more hydrogen bond interactions and tighter coordination with Mn^2+^ stabilize the YdiU-UTP structure, thus leading to a preference of YdiU for UTP over ATP in the presence of Mn^2+^.

### Mechanism for Substrate Recognition of YdiU-mediated UMPylation

Our *in vivo* data suggested that UMPylation occurred on tyrosine or histidine residues that are surrounded with negatively charged and hydrophobic residues. To further explore the mechanism underlying substrate recognition, the peptide from DnaK identified to be UMPylated was synthesized. The interaction between YdiU and this peptide was detected by BLI in the presence and absence of nucleotides and Mn^2+^. Interestingly, no obvious binding signal was detected in the absence of nucleotides (Fig.4I). However, the pre-incubation of YdiU with UTP or ATP significantly enhanced the affinity of YdiU for the DnaK peptide substrate, with a bigger effect of UTP than that of ATP to promote substrate-binding (Fig.4I). To further investigate the mechanism of substrate recognition, four peptides including the one from DnaK were separately docked into the YdiU-UTP-Mn^2+^ model near the active center using ZDCOK software. All of the four polypeptides were reasonably docked to the active site, appropriately matching the surface of YdiU molecule with high ZDOCK scores (Fig.4L, Supplementary Fig.S6). Structural analysis revealed that substrate peptides interact with adjacent residues of YdiU through hydrogen bonds and hydrophobic forces. Remarkably, Arg116 and Asp246 of YdiU were involved in interactions with all four targets, suggesting a universal binding mode for all substrates of YdiU (Supplementary Fig.S6, Fig.S7). Arg116 interacts with negatively charged residues of the substrate through electrovalent bonds, while Asp246 forms a hydrogen bond with the Tyr or His residue that is predicted to be UMPylated. Additionally, Phe76, Met117, and the pyrimidine ring of UTP form a hydrophobic area that can interact with the hydrophobic region of the substrate. This identified substrate recognition feature was completely conserved in different species, suggesting similar substrate selectivity for orthologs of YdiU (Supplementary Fig.S5). Finally, a model for YdiU-mediated UMPyaltion was established: i) Before UTP binding, the essential residue Asp 256 is blocked by Arg170 and Arg 177; ii) UTP and divalent metal bind to the active site through a series of interactions; iii) The hydrophobic region of the substrate is recognized by the hydrophobic pocket, consisting of Phe76, Met117 and UTP. The negatively charged residue near the UMPylation site coordinates with Arg116; and iv) Asp246 activates the substrate for nucleophilic attack, the UTP is hydrolyzed, and the UMP moiety of UTP transfers to the Tyr or His residue of the substrate (Fig.4M, Supplementary Fig.S8).

### Self-AMPylation in Ser478 Inhibits the UMPylation Activity of YdiU

Sreelatha *etal* mapped the major self-AMPylation sites of *E.coli* YdiU to its C-terminal region, 471aa-478aa(Sreelatha et al., 2018). In the structures of YdiU, however, electron density corresponding to C-terminal 468aa-478aa could not be observed suggesting the disorganization of this region in crystals. Sreelatha *etal*. reported an intramolecular disulfide bridge between residues Cys272 and Cys476. To reveal the mechanism of self-AMPylation, the C-terminal peptide of YdiU, WGKRLEVSCSS, was docked onto the cleft between Asp467 and Cys272 of the YdiU-ATP-Mg^2+^ structure. The C-terminal peptide can be positioned near the C-terminal of YdiU well matching the protein surface (Fig.5A). The N-terminal of the peptide is close to the C-terminal of YdiU and Cys272 and Cys476 are close to each other, further supporting the rationality of the molecular docking results. The final residue, Ser478, is predicted to be located outside the active site with a 5.1 Å distance from the α-phosphate of ATP suggesting this might be the major self-AMPylated site of YdiU (Fig.5A). Notably, the C-terminal residues are strictly conserved from bacteria to human, suggesting a uniform mechanism for self-AMPylation (Fig.5B). The exact mechanism of self-AMPylation was further investigated by mutagenesis. Compared with YdiU full-length protein, YdiU^475^ cannot be AMPylated, but YdiU^477^ was able to be AMPylated, suggesting Ser478 and Ser477 can both act as the AMPylated site. Adding one or more Alanine residues to the C-terminal end did not affect AMPylation on Ser478, suggesting the carboxyl group at the C-terminus may not provide a key role (Fig.5C). The mutation of Asp246 completely abolished self-AMPylation, suggesting that Asp246 might be the key catalytic element for self-AMPylation as well as for UMPylation.

**Fig. 5.**
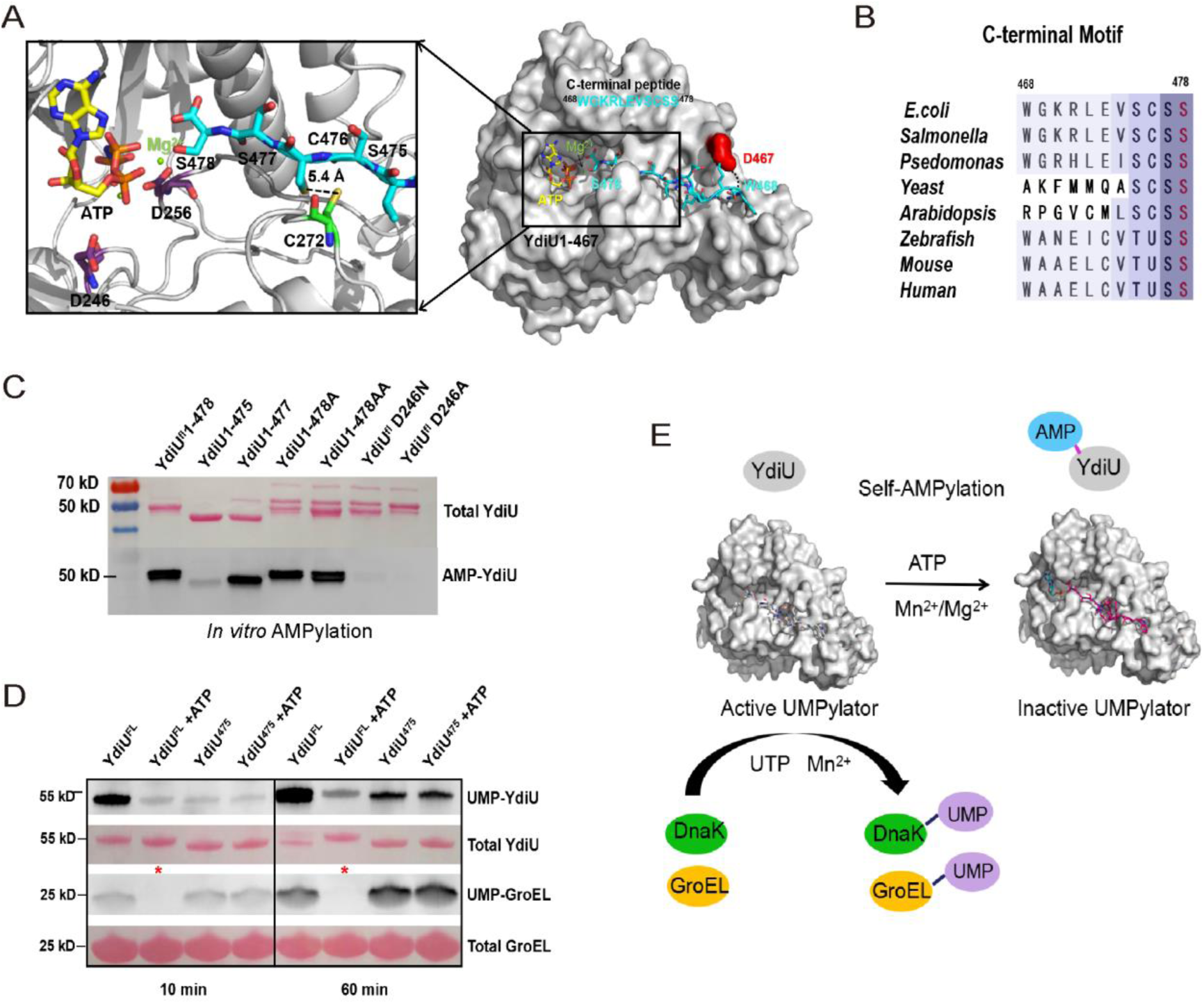
Self-AMPylation inhibits UMPylation activity of YdiU. (A) The C-terminal peptide ^468^WGKRLEVSCSS^478^ was docked into the YdiU-AMPPNP structure. Overall structure is shown in cartoon or surface mode. The peptides and essential residues are shown in stick mode. (B) Sequence alignment of the C-terminal ten residues in YdiU homologues. (C) *In vitro* AMPylation of YdiU and variations performed with biotin-17-ATP and Mg^2+^, as detected by streptavidin HRP blot. Total proteins were visualized by Ponceau S staining. (D) YdiU^fl^ and YdiU^475^ were AMPylated *in vitro* and purified. The UMPylation activities to modify GroEL were detected using biotin-based avidin blotting. Untreated YdiU^fl^ and YdiU^475^ were used as control. Total proteins were visualized by Ponceau S staining. (E) The regulatory model that self-AMPylation inhibits UMPylated activity of YdiU.

To further investigate the effect of self-AMPylation on YdiU-mediated UMPylation, a biotin-based assay was performed. Firstly, YdiU^fl^ or YdiU^475^ was incubated with ATP and Mg^2+^ to generate the Ser478 AMPylated YdiU^fl^ (YdiU^475^ which can not be AMPylated was used as a control). The reaction products were purified and their activities to UMPylate GroEL were detected. The AMPylated YdiU^fl^ lost all the ability to UMPylate GroEL, but the YdiU^475^ that was first incubated with ATP showed similar activity to the non-AMPylated YdiU^475^ suggesting self-AMPylation of YdiU on Ser478 significantly inhibited the UMPylation activity of YdiU. Thus, the UMPylation of substrates mediated by YdiU might be modulated through self-AMPylation in response to the cellular level of ATP (Fig.5E).

### UMPylation Prevents Chaperones from Binding Substrates or Co-factors

As described above, all major bacterial chaperones (GroEL, DnaK, HtpG, and ClpB) were UMPylated by YdiU *in vivo*. To determine how this UMPylation modulates the function of chaperones, the positions of UMPylation were analyzed in depth. The identified UMPylated sites of chaperones were all located in close proximity to the regions responsible for substrate/cofactor binding or self-assembly. Residues Tyr199 and Tyr203 of GroEL are near the hydrophobic pocket and facilitate both substrate and co-factor GroES binding (Fig.6A). As reported previously, mutagenesis of Tyr199 and Tyr203 completely abolished substrate binding(Buckle et al., 1997; Fenton et al., 1994). Tyr199 and Tyr203 of GroEL were further confirmed as the major UMPylation sites by *in vitro* UMPylation assay with GroEL variants in which these two residues were mutated (Fig.6B). Next, the interactions between non-UMPylated GroEL or UMPylated GroEL and human ornithine carbamoyltransferase (OTC, a well-established substrate of GroEL) were monitored using an improved pull-down assay (Supplementary Fig.S9). The results indicated that UMPylation strongly inhibited the GroEL-substrate interaction (Fig.6C), implying that UMPylation can inhibit chaperone function. The UMPylation site of HtpG is also located near the substrate-binding cavity, suggesting that UMPylation of HtpG could affect interaction with substrates (Supplementary Fig.S10A)(Shiau et al., 2006). DnaK requires two cofactors, DnaJ and GrpE, to achieve its chaperone function(Harrison et al., 1997; Liberek et al., 1991). Interestingly, analysis identified the modification site of DnaK as Tyr285, which is located at the interface between DnaK and GrpE (Fig.6D). Biolayer interferometry assays were next performed and further confirmed that UMPylation interfered with the interaction between DnaK and GrpE (Fig.6E and Supplementary Fig.S11). The activity of ClpB requires assembly into a hexamer(Lee et al., 2003). UMPylated residue Tyr812 of ClpB is positioned at the interface between monomers, suggesting that UMPylation of ClpB might influence hexamer assembly and therefore alter ClpB function (Supplementary Fig.S10B). Overall, the UMPylation by YdiU targets residues in these chaperones that could interfere with their function.

**Fig. 6.**
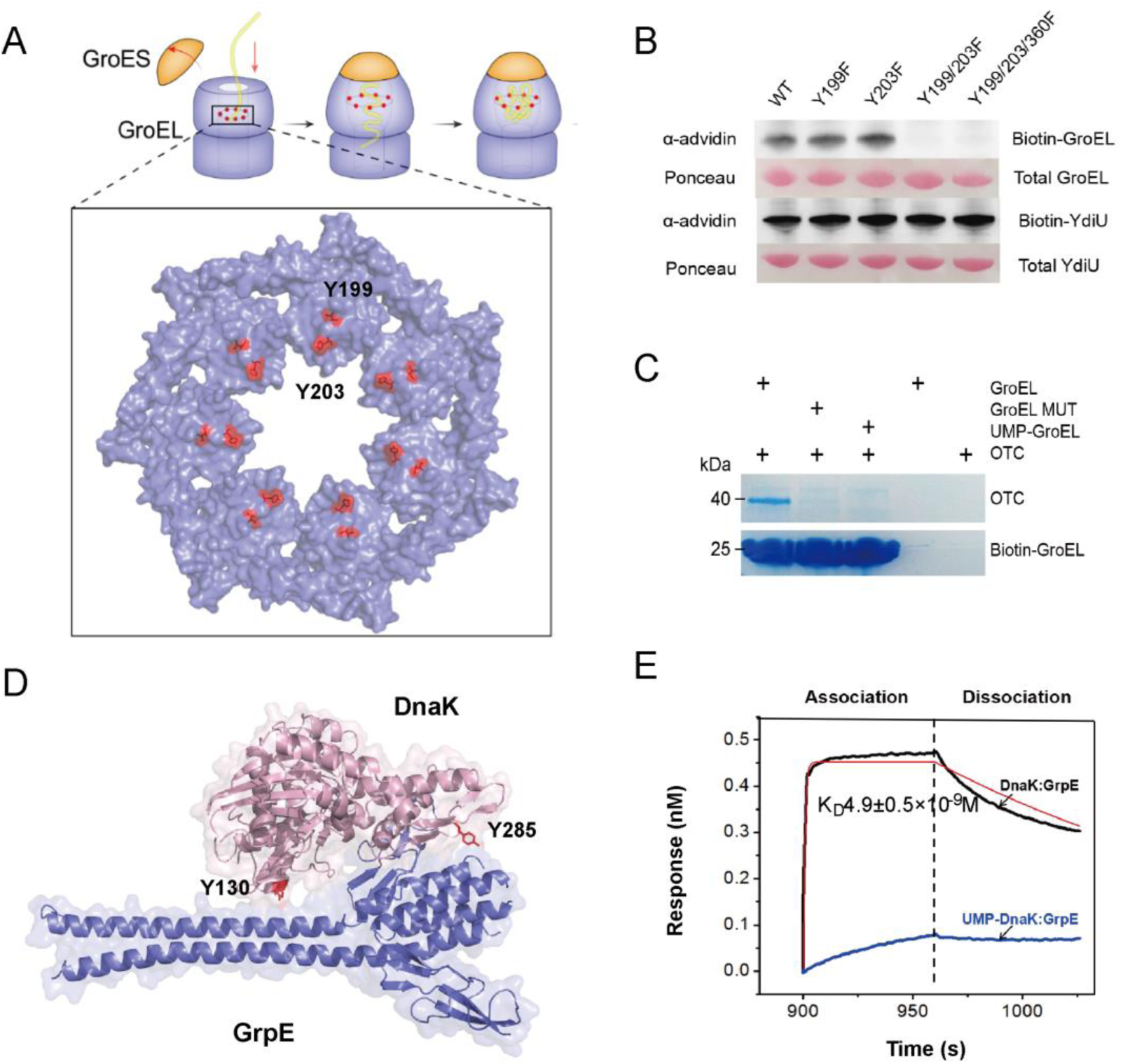
UMPylation of chaperones prevents their binding substrates or co-factors. (A) Structural presentation shows that Y199 and Y203 of GroEL are located in the hydrophobic pore responsible for substrate binding. Both Y199 and Y203 are highlighted as red sticks. PDB code:2CGT. (B) The Y199F/Y203F double mutant could not be UMPylated by YdiU, suggesting that Y199 and Y203 are the major *in vitro* UMPylated sites of GroEL. The UMPylation reactions were performed with Biotin-16-UTP and MnCl_2_ followed by avidin blotting. Total proteins were visualized by Ponceau S staining. (C) The UMPylation of GroEL impairs substrates binding. The interaction between UMP-GroEL and substrate OTC was detected by an improved streptavidin pull-down assay, described in more detail in Supplementary Fig.S9. (D) Y285 and Y130 are located in the interaction surface between DnaK and GrpE. Y285 and Y130 are highlighted as red sticks. PDB code: 1DKG. (E) UMPylation of DnaK inhibits DnaK from binding its co-factor GrpE. The affinities of native DnaK or UMP-DnaK and GrpE were measured by BLI, see also Supplementary Fig.S11.

### YdiU Prevents Stress-induced ATP depletion

We next investigated the role of YdiU in stress response to heat shock. Four *Salmonella* strains were tested: wild-type, the *ydiU* knockout strain (ΔYdiU), or the *ydiU* knockout strain expressing either YdiU (pYdiU) or inactive YdiU mutant D256A (pYdiU D256A). The strains were treated at 55°C for 2 min and their survival rates were determined. Consistent with previous observation (Fig.1G), a remarkable decrease in the survival of ΔYdiU was observed, and this survival defect was fully rescued by expression of YdiU but not the inactive mutant (Fig.7A). The above data suggest that the resistance to heat stress mediated by YdiU depends on the UMPylation activity of YdiU. The UMPylation of chaperones by YdiU can reduce their activities. However, the electron microscopy observation of recovering cells after heat treatment (55°C, 2 min) showed no obvious difference in protein aggregation in the ΔYdiU and WT strain (Supplementary Fig.S12). The expression of YdiU and DnaK after heat shock were analyzed. *Salmonella* elevates the expression of DnaK less than 5 min after heat exposure, while YdiU expression is induced 10 min after heat, and only reaches its highest level 45 min after heat treatment (Fig.7B). Thus, the UMPylation of chaperones will happen once YdiU has been induced, around 30 min post-heat-treatment, which may be after chaperones have already acted to refold misfolded polypeptides that resulted from the heat. YdiU may remove excess or inoperative chaperones after the acute recovery phase. Recovery from heat injury consumes lots of ATP. Under ATP-limited condition, the induced YdiU will inactivate chaperones through UMPylation. *Salmonella* might benefit from this process for improved survival under severe stress condition by effectively and rapidly turning off the ATP consumption of chaperones in response to low ATP level. To check this role of YdiU in energy management under stress condition, the intracellular levels of ATP were monitored in WT and ΔYdiU *Salmonella* during heat recovery. Notably, the intracellular concentration of ATP in ΔYdiU continuously decreased upon heat treatment, while that of the WT *Salmonella* initially declined and then increased upon YdiU expression. Two hours post-heat treatment, the ATP level of WT *Salmonella* recovered to the level before heat, however, that in ΔYdiU still remained at a very low level (Fig.7C). The heat-induced ATP depletion phenotype of ΔYdiU was improved by the expression of functional YdiU but not by expression of the inactive mutant (Supplementary Fig.S13). The above data demonstrate a regulatory role of YdiU in energy utilization, which might dramatically improve the survivability of *Salmonella* under severe stress. Given that self-AMPylation inhibits chaperone-UMPylation, YdiU may act as an energy meter to switch off chaperone function in response to the level of ATP.

**Fig. 7.**
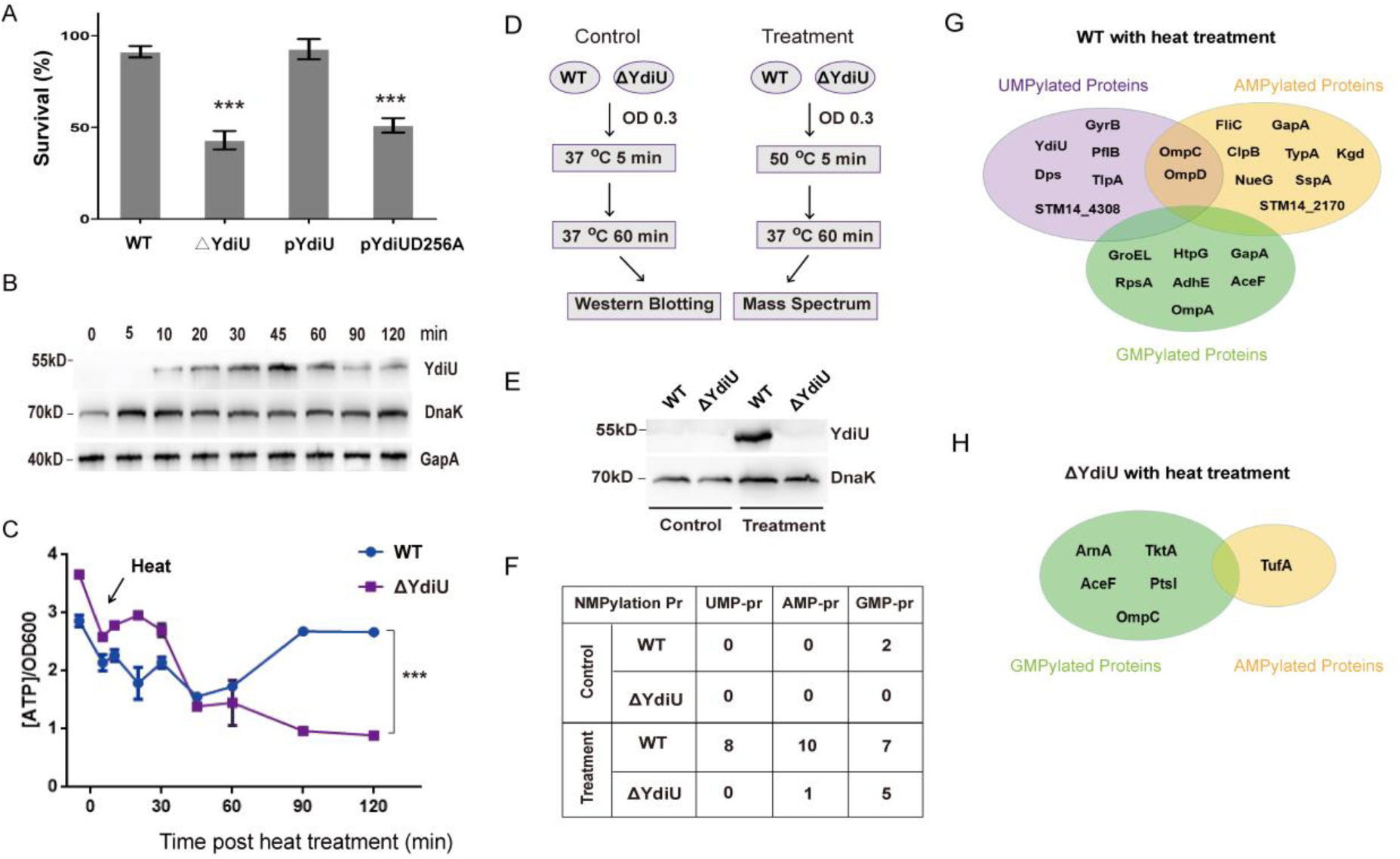
YdiU prevents stress-induced ATP depletion of *Salmonella* through UMPylation. **(A)** Survival ratios of indicated *Salmonella* strains were measured following treatment at 55°C for 2 min. The statistical significance is indicated by ***P < 0.001 as compared with wild type strain. (B) At the indicated time points after heat treatment (55°C, 2min), the expression levels of YdiU and DnaK of WT Salmonella were quantified by western blot. GapA (also known as GADPH) was used as a loading control. (C) ATP levels of WT and ΔYdiU *Salmonella* during recovery from heat injury. All experiments were performed in triplicate and the mean values and error bar are presented. The average amount of intracellular ATP was obtained by calculating the ratio of total ATP and OD600. Statistical significance is indicated by ***P < 0.001 using a t-test. Also see Supplementary Fig.S13. (D) Schematic illustrating the preparation of the heat-treated and control samples. (E) The YdiU and DnaK expression levels in treatments and controls were detected by western blot analysis. (F) Shot-gun mass spectrometry-based proteomics were performed using treatments and controls. The number of identified nucleotide-mediated PTMs (UMPylation, AMPylation and GMPylation) were shown. (G) The modified proteins identified in heat-treated WT *Samonella* were classified by UMPylation, AMPylation and GMPylation. (H) The modified proteins identified in heat-treated ΔYdiU *Samonella* are classified by UMPylation, AMPylation and GMPylation.

### UMPylation Occurs in Response to Stress in a YdiU-dependent Manner

To further investigate the relationship between YdiU, UMPylation, and stress signaling *in vivo*, further experiments were performed with WT and ΔYdiU *Salmonella* strain before or after heat treatment (Fig.7D). As shown earlier and presented in Fig.1D, YdiU is highly expressed in heat-treated WT *Salmonella* but not in the untreated culture (Fig.7E). Mass spectrometry-based proteomics were performed on these treated strains. Twenty-three proteins with NMPylation modifications (AMPylation, UMPylation, and GMPylation) were identified in heat-treated WT *Salmonella,* but only two of these modifications were identified in the untreated WT *Salmonella* suggesting NMPylation might be induced by heat stress (Fig.7F-H, Supplementary Table S4). No UMPylation of proteins was detected in the ΔYdiU strain, suggesting UMPylation occurred in a YdiU-dependent manner (Fig.7F, Supplementary Table S4). Remarkably, both AMPylation and GMPylation of proteins were detected in heat-treated ΔYdiU, implying other unknown enzyme(s) might mediate AMPylation and GMPylation in *Salmonella*.

## DISCUSSION

The attachment of UMP to proteins has been rarely reported. The sole identified prokaryotic UMPylator is GlnD, from the glutamine synthetase system, which acts to UMPylate its single substrate GlnB to modulate the synthesis rate of glutamine(Adler et al., 1975). AvrAC, an effector from *Xanthomonas*, has been reported to UMPylate *Arabidopsis* BIK1 and RIPK proteins, thereby inhibiting host immune signaling(Feng et al., 2012). Intriguingly, AvrAC employs a Fic domain to achieve UMPylation activity, which is an unusual activity for this domain, as most other Fic domains modify targets with AMP(Casey and Orth, 2018; Worby et al., 2009; Yarbrough et al., 2009). YdiU shares no sequence identity with either GlnD or AvrAC, but our results show that in *Salmonella*, this protein UMPylates forty-six bacterial substrates. YdiU-dependent UMPylation occurs primarily on the tyrosine and histidine residues. To our knowledge, YdiU is the first known enzyme to catalyze UMPylation of histidine. Histidine and tyrosine residues play significant roles in enzymatic catalysis and protein-protein interactions, and the linking of an UMP moiety to those residues may have a significant effect on protein function. The YdiU mutant strain showed an obvious defect in resisting with multiple stress conditions compared with the wild type strain, and we speculate that YdiU may act as a global regulator to modulate stress signaling of bacteria through the UMPylation process. Our research further revealed an essential role of YdiU and UMPylation in modulation of the chaperone network and the thermo-resistance of bacteria. Further study should explore these activities in higher eukaryotes.

Our data demonstrate that the UMPylated activity of YdiU might be regulated by many factors in response to multiple cellular signaling. i) Mn^2+^ plays an important role in YdiU-mediated UMPylation indicating a regulatory mechanism to sense the intracellular level of metal ions. ii) Self-AMPylation entirely abolished substrate-UMPylation of YdiU suggesting that UMPylation is modulated in response to the cellular level of ATP or the ATP/UTP ratio. iii) The UMPylated activity of YdiU might be affected by the redox state in cells. Consistent with this, the self-AMPylation of YdiU in Ser478 is regulated by formation of a disulfide bridge between Cys272 and Cys476. High eukaryotic homologs of YdiU include a redox-active selenoprotein located in mitochondria(Han, 2014). The AMPylated substrates of YdiU were identified to be involved in redox homeostasis confirming a relationship between YdiU and redox process(Sreelatha et al., 2018). iv) The UMPylation level of chaperones might be dynamically adjusted by the ratio of misfolded protein and the level of YdiU in cells. The UMPyaltion sites of chaperones are located in key regions for substrate/cofactor binding suggesting the possibility that YdiU could modify chaperones in a substrate-unbound state, or that YdiU might compete with substrate for chaperone binding sites. Our data demonstrate that UTP and Mn^2+^-binding converts YdiU to an unstable state that tends to aggregate. Structural analysis showed that UTP and Mn-loaded YdiU form a hydrophobic center consisting of Met117, Phe76 and UTP to recognize the hydrophobic region of substrates. Since hydrophobic residues act as the core-element of chaperone-binding sites, we suspect that the UTP and Mn-loaded YdiU could mimic chaperone clients to recruit chaperones and then UMPylate them. YdiU might compete with misfolding proteins for chaperone binding sites, as a result, the ratio of YdiU and misfolding proteins generated by stress directly governs the level of UMPylated chaperones. Very recently, the bacterial effector HopBF1 was reported to mimic the target of its host HSP90 to phosphate HSP90(Lopez, 2019), indicating this “betrayal-like” mechanism might exist widely to regulate chaperone function through post-translational modification. The *in vivo* regulation of YdiU-mediated UMPylation may be a complex process combining overall effects of these various elements.

Recently, Sreelatha *et al* reported the crystal structure of *P. syringae* YdiU-ATP and further identified YdiU as an AMPylator that can modify itself and two proteins involved in redox homeostasis(Sreelatha et al., 2018). Consistent with that finding, we found that YdiU exhibits *in vitro* self-AMPylation activity. However, mass spectrometric assays of YdiU-expressing *Salmonella* failed to detect AMPylation targets of YdiU, although this method was successfully used to detect dozens of targets with UMPylation. Previously, a mutual regulation between AMPylation and UMPylation was established for the regulation of glutamine synthetase (GS) by Earl Stadtman(Stadtman, 2001). Glutamine synthetase adenylyltransferase (GlnB) AMPylates GS, thereby inactivating this enzyme, and when GlnB is modified with UMP by GlnD, UMPylated GlnB can catalyze the deAMPylation of GS to re-activate GS. In that example, UMPylation and AMPylation work together to regulate the homeostasis of glutamate synthesis. With regard to YdiU, self-AMPylation may act to regulate its UMPylated activity. Self-AMPylation by YdiU serves as a regulatory mechanism of enzymatic activity in response to ATP level, but UMPylation may be the real physiological function of YdiU to modify and regulate substrates. Another possibility is that UMPylation or AMPylation by YdiU function in different pathways in response to different signals. Whether YdiU regulates cell activity through both AMPylation and UMPylation processes requires further investigation.

Before UMPylation was identified and characterized in this study, phosphorylation and glutathionylation processes were reported to regulate the activities of bacterial chaperones. DnaK and GroEL can autophosphorylate in response to high temperature, therefore acting to enhance their ATPase activity and increase their affinity for unfolded proteins(McCarty and Walker, 1991; Sherman and Goldberg, 1994). Glutathionylation of DnaK, however, acts to reduce its activity by changing both its secondary structure and tertiary conformation(Zhang et al., 2016). In higher eukaryotes, the activities of endoplasmic reticulum (ER)-localized Hsp70 chaperone BiP and cytosolic chaperone Ssa2 are regulated by AMPylation, which is mediated by Fic domain proteins in response to unfolded protein stress or heat stress(Ham et al., 2014; Sanyal et al., 2015; Truttmann et al., 2017). Recently, aggregation and toxicity of neurodegenerative disease-associated polypeptides were found to be modulated by the AMPylation of chaperone in C.elegans(Truttmann et al., 2018). This is the first report of the nucleotidylation of chaperones in prokaryotes. Our data show that UMPylation of chaperones by YdiU inhibits their activities. The biological significance of YdiU-mediated UMPylation of chaperones might be related to the conservation of energy during stress response. The recovery from stress is an energy-consuming process for bacteria, requiring dozens to thousands of ATPs to refold a single misfolded protein(Diamant and Goloubinoff, 1998; Sharma et al., 2010). Chaperones, such as DnaK, can effectively repair small protein aggregates, but they may be unable to refold large protein aggregates containing too many chaperone-binding sites even with the consumption of lots of ATP(Diamant, 2000). Stresses frequently lead to the depletion of cellular ATP. When ATP is limited, turning off chaperone activities by YdiU through the process of UMPylation would allow bacteria to maintain sufficient energy for their basic life activities and protect bacteria from ATP depletion during recovery. The observation of the sharp decline of ATP in the ΔYdiU strain during heat recovery but not in wild type *Salmonella* supports this hypothesis (Fig.7C). Under stress condition, bacteria need to estimate if they have the ability to repair all the misfolded proteins. When chaperones have done most of their job, it is better to turn off chaperone function to avoid excess utilization of ATP, and YdiU can function in this process.

On the basis of the above data, we propose a model in which YdiU modulates chaperone activity in response to both stress condition and ATP level (Supplementary Fig.S14). First, the expression of YdiU is induced by stress. Chaperones repair the misfolded proteins using ATP. When the intracellular concentration of ATP is in a high level, YdiU is self-AMPylated and inactive. In the later stage of recovery, most of misfolded proteins were refolded by chaperones, which consume lots of ATP. Under ATP-limited condition, YdiU is activated as an UMPylator. Then, YdiU will UMPylate and inactive chaperones to prevent stress-induced ATP depletion.

It is very interesting that GMPylation of proteins is observed in *Salmonella*. To our knowledge, this is the first report that the GMPylation of protein exists in bacterial cells. More importantly, the GMPylated sites present a similar pattern, in a Lys followed by an Asp, making these sites very likely to be catalyzed by the same enzyme. This activity cannot be from YdiU because GMPylated proteins were also identified in heat-treated ΔYdiU, so the enzyme that catalyzes the GMPylation of proteins and its role in bacteria are valuable subjects for further investigation.

In conclusion, our studies illuminate new biological functions of the YdiU domain family, highlight the importance of UMPylation in bacterial signal transduction, and reveal a potential mechanism by which bacteria can adapt to a stressful environment.

## Supporting information

supplemental informations

## ACKNOWLEDGEMENT

We acknowledge the staffs at beamline BL17Ul and BL19Ul of the Shanghai Synchrotron Radiation facility for supporting data collection of protein crystals. We acknowledge Dr. Jiawei Wu (GE Healthcare) for his help in SPR experiments and Dr. Scott Zhou (Unchained Labs) in UNcle experiments.

## Author contributions

B.L. and L.G. designed the study; YL.Y., YY.Y., Y.M. and H.D. purified proteins; YY.Y., C.L., YL.Y. and H.L. performed *in vitro* UMPylation experiments; F.Z. constructed *ydiU* knock-out strain; N.S., Y.W., P.L. and X.L. performed *in vivo* experiments; B.L., Z.Y. and L.G. performed structural experiments; YL.Y., W.W. and H.J. performed BLI assay; YL.Y., Q.W. and J.D. performed SPR assay; B.L., YY.Y., C.L. and YL.Y. analyzed data; B.L. wrote the manuscript.

## DATA AVAILABILITY

The X-ray structures (coordinates and structure factor files) of YdiU have been submitted to PDB under accession number 6INY (YdiU-AMPPNP), 6III (YdiU-AMP-imidodiphosphonic acid) and 6K20 (Apo-YdiU). Mass spectrometry based data were submitted to ProteomeXchange under accession number PXD013825.

## STAR METHODS

### Bacterial strains and culture media

The bacterial strains used in this study are listed in Supplementary Table S5. For the experiments shown in Fig.1, *S. typhimurium* ATCC14028 culture was grown overnight and subcultured into corresponding medium. When the OD600 reached 0.4-0.5, bacterial cells were harvested for subsequent functional study. LB medium and M9 minimal medium were prepared as described previously(Wada et al., 2011). H_2_O_2_-stress medium (H_2_O_2_) was LB medium supplemented with H_2_O_2_ to a final concentration of 1 mM. Iron-limited medium (Fe^−^) was LB medium supplemented with 2,2-dipyridyl to a final concentration of 0.5 mM. Antibiotic-stress medium (AMP) was LB medium supplemented with 0.5 μg/mL Ampicillin. High osmotic stress medium were prepared by adding 300 mM NaCl (NaCl) or 20% w/v Glucose (Glu), respectively. Acid-stress medium (H^+^) was LB medium supplemented with 0.5 μl/mL acetic acid (∼pH 5.2). Envelop-stress medium (Indole) was LB medium adding indole to a final concentration of 2 mM. For the experiments shown in Fig. S1, YdiU was induced using LB medium supplemented with 0-0.2% L-arabinose.

For the experiments shown in Fig.1B, 1C and 7E, indicated strains were grown overnight in 10 mL LB medium. 50 μl above culture was subcultured into 5 mL LB medium. When the OD600 reached 0.3-0.4, strains were treated as indicated time at indicated temperature. After recovery at 37°C for 30 min (Fig.1B and 1C) or 60 min (7E), bacterial cells were harvested for subsequent functional study.

### RNA isolation and real-time quantitative PCR

Total RNA was isolated using the RNAprep Pure Bacteria Kit (TIANGEN). The reverse transcription reactions were performed using HiScript II Q-RT SuperMix (Nanjing Vazyme BioTech Co.,Ltd.) according to manufacturer’s instructions and incubated at 42° for 60 min, followed by 10 min at 70°. The quantitative RT-PCR reactions were performed on an Applied Biosystems 7500 Sequence Detection system (Applied Biosystems, Foster, CA, USA) using ChamQ SYBR Color qPCR Master Mix (Nanjing Vazyme BioTech Co.,Ltd.). GADPH was used as an internal reference.

### Antibody preparation

Antibody against YdiU and GapA (reference) were prepared by Dia-An Biotech, Inc, in Wuhan, China. 2mg purified protein was mixed with FCA (Freund’s complete adjuvant) or FIA (Freund’s incomplete adjuvant), followed by injecting into two Japanese White rabbits four times. FCA is only used in the first injection, while FIA for the rest injections. The 2nd injection is at 28th day since the first injection. And for other injections, 2 weeks intervals were needed between each injection. 3 days after the 4th injection, antiserum titer were tested by ELISA. The rabbits with higher titer was killed and bleeded at the 64^th^ day since the first injection. The affinity column was made by coupling 1mg purified protein to CNBr-activated Sepharose 4B from GE. The antiserum was applied onto the column, the specific antibody were eluted with the Glycine HCl buffer at pH 2.5.

### Western blot

Bacterial cells were lysed using 4×SDS-PAGE loading dye followed by heating at 95°C for 10 min before SDS-PAGE. Total proteins on gels were transferred to nitrocellulose membranes at 250 mA for 2 h in transfer buffer (96 mM glycine, 12.5 mM Tris, and 10% methanol). The membranes were blocked in 5% milk in PBS-0.1% and Tween 20 (PBST) at 37°C for 1h, followed by incubation with polyclonal antibody to YdiU (this study) diluted 1:2000, GapA (this study) diluted 1:5000, DnaK (Abcam) or GroEL (Novusbio) diluted 1:10000 in PBST overnight at 4°C. The membranes were incubated with HRP-Conjugated Goat anti Rabbit or Mouse IgG (h+l) (Abcam) diluted 1:10000 in PBST at 37°C for one hour after three washes with PBST. The membranes were then washed three times and then detected by chemiluminescence.

### Construction of *ydiU* knock-out strain

S. *typhimurium* ATCC14028 strain was purchased from American Type Culture Collection. The *ydiU* gene knockout strain was constructed using the lambda Red recombinase system as described previously(Datsenko and Wanner, 2000). Briefly, the chloramphenicol resistance-FRT cassette was amplified from pKD3 by PCR with primers including the 5’and 3’ flanking regions of the *ydiU* gene. The amplified product was transformed into *S. typhimurium* ATCC14028 strain containing the pKD46 plasmid, and then recombined colonies were selected on LB agar plates containing 25 μg/ml chloromycetin. PCR was then performed with another two primers to confirm the deletion of the *ydiU* gene. The pCP20 plasmid was then used to remove the chloramphenicol resistance gene. The *ydiU* knock-out strain of *E.coli* BL21 (DE3) was constructed as described above.

### Plasmid construction

Plasmids used in this study are listed in Supplementary Table S5. For *in vivo* study, *Salmonella* YdiU 1-475aa and *Salmonella* YdiU 1-475aa D256A was cloned into the pBAD24 vector. For biochemical study, YdiU full-length protein, YdiU1-477aa, YdiU1-478+A, YdiU1-478+AA, YdiU1-475aa, GroEL191-376aa, DnaK, HtpG 1-624aa, ClpB, GrpE, GrxA genes were amplified from *E.coli* str. K12 substr. MG1655 genomic DNA and cloned into an expression vector, pGL01, a modified vector based on pET15b with a PPase cleavage site to remove the His tag. Mutants of YdiU (Y71A, K107A, D119A, E130A, N244A, D246A, D246N, N247A, D256A), GroEL191-376aa (Y199F, Y203F, Y199F-Y203F, Y199F-Y203F-Y360F) were constructed using a two-step PCR strategy and separately cloned into pGL01.

### Sample preparation for mass spectrometry

For the experiments shown in Fig.2, the recombinant plasmid encoding *Salmonella* YdiU1-475aa was transformed into the *ydiU* knockout strain to generate the YdiU-expressed strain (pYdiU) while empty pBAD24 was transformed into the *ydiU* knockout mutant as a negative control (ΔYdiU). Both ΔYdiU and pYdiU strains were grown overnight in LB medium with 100 μg/ml Ampicillin to select for the plasmids, and then subcultured into 10 mL LB medium with 100 μg/ml Ampicillin. When the OD600 reached 0.4-0.6, 0.1% L-arabinose was added to induce YdiU expression. Bacterial cells were harvested 4 hours later.

For samples shown in Fig.7, wild type and ΔYdiU *Salmonella* were grown overnight in 10 mL LB medium. Then 50μl above culture was subcultured into 5 mL LB medium. When the OD600 reached 0.3, strains were treated with 50 °C for 5min, recovered at 37°C for 60 min and harvested. Strains without heat treatment are used as controls.

Samples are lysed using a buffer containing 4 % SDS, 100mM Tris-HCl pH7.6, 1mM DTT and lysozyme. After centrifugation at 14000 rpm for 40 min, the supernatant was removed and quantified with the BCA Protein Assay Kit (Bio-Rad, USA). Samples containing 200 μg of proteins were purified by ultrafiltration, incubated with iodoacetamide to block reduced cysteine residues, and then digested with 4 μg trypsin at 37 °C overnight. Peptides of each sample were desalted on C18 Cartridges and concentrated by vacuum centrifugation and reconstituted in 40 µl of 0.1% (v/v) formic acid.

### Mass spectrometry

Peptides were fractionated by SCX chromatography using the AKTA Purifier system (GE Healthcare) and injected for nanoLC-MS/MS analysis. LC-MS/MS analysis was performed on a Q Exactive mass spectrometer (Thermo Scientific) that was coupled to Easy nLC (Proxeon Biosystems, now Thermo Fisher Scientific) for 240 min in positive ion mode. MS/MS spectra were automatically searched against the Uniprot_Salmonella_typhimurium_99287 database using the ProteinPilot software 4.5 (AB Sciex) to identify the protein fragments and the protein identification results were obtained. UMPylated peptides, AMPylated peptides and GMPylated peptides were identified by searching for the modification ‘PhosphoUridine’, ‘Phosphoadenosine’, ‘Phosphoguanosine’, respectively.

### Protein expression and purification

Proteins were expressed in *E. coli* BL21(DE3). When the OD600 reached 0.4-0.6, cultures were cooled to 16°C and induced overnight by 0.01-0.1 mM IPTG. Harvested cells were lysed by sonication. Proteins were purified by Ni^2+^-NTA affinity column. The His-tag of the proteins was removed by PPase treatment(Li et al., 2017). Then proteins were then concentrated and purified by Superdex 200 chromatography. Selenomethionine-labeled YdiU was expressed in *E. coli* BL21(DE3) using the methionine biosynthesis inhibition method(Hendrickson et al., 1990).

### Crystallization and structure determination

YdiU^475^ was concentrated to 10 mg/ml. Crystals of Apo-YdiU were grown by hanging drop vapour diffusion in a buffer containing 10% PEG200; 0.1M Bis-Tris propane pH9.0; 18% PEG8,000. To obtain YdiU-AMPPNP-Mg^2+^, 10mM AMPPNP and 10 mM MgCl_2_ were added to YdiU and crystals of YdiU-AMPPNP appeared in a buffer containing 2.8 M Sodium acetate pH7.0 and 0.1 M Bis-Tris propane pH7.0. To obtain YdiU-AMPNPP-Mn^2+^, 10mM AMPNPP and 1 mM MnCl_2_ were added to YdiU. The crystals obtained in the buffer containing 0.1 M Sodium Cacodylate pH6.3, 0.2 M Calcium Acetate and 18% PEG8,000. Then, 10mM UMPNPP and 1 mM MnCl_2_ were added to YdiU to generate YdiU-UTP-Mn^2+^ complex, however, protein precipitated after this operation. 2mM UMPNPP and 0.5 mM MnCl_2_ were used as well, no improvement is observed.

Diffraction data of the above crystals were collected at the Shanghai Synchrotron Radiation facility (SSRF) using beamlines BL17u1 and BL19ul. The data sets were processed using the HKL2000 software suite(Otwinowski and Minor, 1997). The structure of YdiU-AMPPNP-Mg^2+^ was solved by selenium single wavelength anomalous diffraction and those of Apo-YdiU, YdiU-AMP-PPi-Mg^2+^ and YdiU-AMNPP-Mn^2+^ were solved by molecular replacement. The initial model was generated using the program Autosolve(Terwilliger, 2002). The subsequent model was manually built using COOT and refined using PHENIX(Adams et al., 2002; Emsley and Cowtan, 2004). The data collection and structure refinement statistics are summarized in Supplementary Table S6. Structural figures were generated using PyMol (http://www.pymol.org).

### Molecular docking

UTP was docked to the binding site of YdiU using Autodock Tools 1.5.6 performing using a flexible UTP molecule and a rigid YdiU structure(Goodsell et al., 1996). Application of an Amber ff14SB force field was used to obtain the appropriate conformations and binding positions of UTP by energy optimization(Maier et al., 2015). Finally, the structure with the lowest docking energy was selected as the final model for YdiU-UTP. The number of grid points in the XYZ of the grid box was set to 40×40×40, the grid spacing was 0.375 Å, the number of GA run was set to 100, and default settings were used for the remaining parameters. Finally, the structure with the lowest docking energy was improved with energy minimization. This optimization process was carried out in two steps, using a steepest descent method optimization of 2000 steps, followed by optimization of this model by 2000 steps with the conjugate gradient method.

To study the binding of YdiU to polypeptide molecules (Fig.4L, 5A and S6), the rigid-body docking program Zdock was used to predict complex structures of peptides bound to YdiU protein(Pierce, 2011). The version of software ZDOCK3.0.2 was used in this study (http://zdock.umassmed.edu). The ZDOCK score can be used to evaluate the shape complementation, electrostatic potential and energy matching of proteins. The predicted structure with the highest ZDOCK score was used for further optimization. Moreover, Amber 14 force field was used to obtain the appropriate conformations by energy optimization. The optimization process was carried out in two steps of steepest descent method optimization of 5000 steps, followed by additional optimization of the structure by 5000 steps with the conjugate gradient method.

### Molecular dynamics simulation

MD simulation of YdiU-ATP or YdiU-UTP complex was applied using the Amber 18 software package(Case et al., 2005). Amber ff14SB force field was applied, in which the coordination of Mg^2+^ or Mn^2+^ with the surrounding residues was optimized by MCPB(Li and Merz, 2016). The parameters of the protein field were set based on the experimental condition. The simulated temperature was set as 300 K and the pH was set as 7.0. The solutes were solvated in a truncated periodic box with a 1.0 nm distance solute-wall using the TIP3P water model(Price and Brooks, 2004). Prior to MD simulation, the system was optimized by application of the following two steps. First, the solute was constrained, and the water solvent was optimized using 5000 steps with the steepest descent method and 5000 steps with the conjugate gradient method. Second, unrestrained minimization of the whole system was executed with another 5000 steps of steepest descent and 5000 steps of conjugate gradient. The MD simulation process was also carried out in two stages. First, a 100 ps-constrained solute MD simulation was performed in which the system temperature was gradually increased from 0K to 300 K, and then 50 ns unconstrained constant temperature MD simulation was performed. In the simulation process, the SHAKE algorithm was applied to constrain covalent bonds including hydrogen atoms(Coleman et al., 1977). The MD time step was set as 2 fs, with one snapshot was taken every 10 ps (1 ps), for a total of 5,000 conformations in each MD simulation. The MD simulation dynamic process was monitored with the VMD software package(Humphrey et al., 1996).

### *In vitro* UMPylation assay with α^32^P-UTP

For the data shown in Fig.2H, 2 μg of purified YdiU^475^ or YdiU^fl^ was incubated in a 20 μl UMPylation reaction buffer containing 25 mM Tris-HCl (pH7.5), 1 mM DTT, 25 mM MgCl_2_/CaCl_2_/ZnCl_2_/MnCl_2_, and 2 µCi α^32^P-UTP at 30 °C for 30 min. Then products were analyzed using 12% NuPAGE gels and imaged by autoradiograph.

For the experiment shown in Fig.2I, 8 μg of the corresponding chaperones was incubated with or without 1 μg YdiU^475^ in a 20 μl reaction buffer with 10 mM MnCl_2_ and 2 µCi α^32^P-UTP at 30 °C for 30 min, then analyzed using 12% NuPAGE gel and imaged by autoradiograph.

### *In vitro* UMPylation/AMPylation assay with Biotin-16-UTP or Biotin-17-ATP

For the experiment shown in Fig.2J, 8 μg of purified chaperones were incubated with or without 1 μg YdiU^475^ in a 20 μl reaction buffer containing 25 mM Tris-HCl (pH7.5), 1 mM DTT, 100 mM NaCl, 10 mM MnCl_2_, and 500 μM Biotin-16-UTP. For the AMPylation experiment shown in Fig.3E, 2 μg GrxA were incubated with or without 1 μg YdiU^fl^ in a 20 μl reaction buffer containing 25 mM Tris-HCl (pH7.5), 1 mM DTT, 100 mM NaCl, 10 mM MgCl_2_/MnCl_2_, and 500 μM Biotin-17-ATP. The self-AMPylation reaction shown in Fig.5C were performed using 1 μg YdiU or variants with 10 mM Mg^2+^ and 100 μM Biotin-17-ATP. For the experiment shown in Fig.3F and 3G, 1 μg YdiU^fl^ were used to modify 4 μg DnaK in a buffer containing 10 mM MnCl_2_/MgCl_2_ and 100 μM Biotin-16-UTP/Biotin-17-ATP. For the experiment shown in Fig.3H, 1 μg YdiU^FL^ or YdiU^475^ were used to modify 4 μg GroEL in a buffer containing 25 mM Tris-HCl (pH7.5), 1 mM DTT, 100 mM NaCl, 10 mM MnCl_2_, 10 mM MgCl_2_ and 100 μM Biotin-16-UTP or Biotin-17-ATP. For the experiment shown in Fig.4K and Fig.6B, 1 μg native YdiU or variants was incubated with 2 μg native GroEL or variants in the same buffer as Fig.3H. After incubation at 30 °C for 1h, the above reaction products were resolved by 12% NuPAGE gel and transferred to nitrocellulose membranes. The membranes were stained in 0.1% Ponceau S with shaking for 5 min at room temperature, and then washed three times with ddH_2_O. The Streptavidin HRP blot was performed as previously described(Roux et al., 2013). Briefly, the membranes were incubated in BSA blocking buffer (1×PBS containing 1% BSA and 0.2% Triton X-100) with shaking for 30 min at room temperature. Next, the membranes were incubated with streptavidin-HRP at 1:40,000 dilution in BSA blocking buffer overnight at 4°C. The membranes were washed three times with 1×PBS and then blocked with ABS blocking buffer (1×PBS containing 10% adult bovine serum and 1% Triton X-100) with shaking for 5 min at room temperature. Membranes were washed three times with 1×PBS and blots were developed and visualized using a enhanced chemiluminescence (ECL) kit.

For the experiment shown in Fig.5D, AMPylated YdiU^fl^ and YdiU^475^ were product in a reaction buffer containing 25 mM Tris-HCl (pH7.5), 1 mM DTT, 10 mM MgCl_2_ and 10 mM ATP at 30°C for 4 h and purified using desalting column (GE, PD MiniTrap G-25) to remove ATP and Mg^2+^. Then 1 μg AMPylated or non-AMPylated YdiU was incubated with 4 μg purified GroEL in the UMPylation buffer containing 25 mM Tris-HCl (pH7.5), 1 mM DTT, 10 mM MnCl_2_ and 100 μM Biotin-16-UTP at 30°C for 10min or 60 min.

### Microscale Thermophoresis (MST) assay

The binding affinity of the purified YdiU to Mn^2+^ or Mg^2+^ was measured using a Monolith NT.115 (NanoTemper Technologies). YdiU was fluorescently labeled with according to the manufacturer’s procedure. Briefly, the concentration of YdiU was adjusted to 10 μM and then incubated with red fluorescent dye RED-tris-NTA (L018) for 30 min at 25°C in the dark. Next, the labeled YdiU (about 100 nM) was mixed with the same volume of Mn^2+^/Mg^2+^ in 12 serial concentrations. The samples were then loaded into premium capillaries (NanoTemper Technologies) and measured at 25 °C by using 100% excitation power and 40% MST power. Each assay was repeated two times, and data analyses were performed using MO. Affinity Analysis v.2.2.4 software.

### Biolayer interferometry (BLI) assay

Biolayer interferometry experiments were performed using an Octet RED96 instrument (ForteBio) at 25°C. To study the interaction between YdiU and DnaK peptide (Fig.4I), purified YdiU^475^ was incubated with EZ-Link-Biotin (MCR=3:1) at 25°C for 30 min, and then purified by passage through a desalting column (GE, PD MiniTrap G-25). About 100 μg/ml YdiU was loaded onto each Super Streptavidin senor (ForteBio, 18-5057) in PBS buffer. Following a baseline step, the sensors were introduced to HBS buffer (25 mM HEPES pH7.5, 100 mM NaCl_2_) containing 50 μM MnCl_2_ or 50 μM MnCl_2_+25 μM UMPNPP or 50 μM MnCl_2_+AMPNPP for 300 s to perform Mn^2+^ and nucleotide-binding. After equilibration for 130s, the corresponding buffer containing 100 μM peptide was applied for 60 s to measure the binding signal and then the buffer without peptide was used to detect dissociation. A YdiU-loaded sensor in HBS buffer was used as a control. To study the effect of UMPylation on the interaction between DnaK and GrpE (Fig.6E), experiments were performed as described in Fig.S11. Briefly, DnaK proteins were biotinylated either by EZ-Link-Biotin or by biotin-16-UTP and YdiU, and then separately loaded to streptavidin sensors. The sensors were introduced to PBS containing 10 μg/ml GrpE to measure the association response and changed to PBS buffer to measure dissociation. Data were processed by deduction with sensor control.

### Surface plasmon resonance (SPR) assay

The binding affinities between YdiU and Mn^2+^ or a nucleotide analogue were performed by surface plasmon resonance (SPR) on a Biacore T200 instrument (GE Healthcare Life Sciences).

First, purified YdiU^475^ in HBS buffer (25 mM HEPES pH7.5, 100 mM NaCl_2_) was diluted with 10 mM sodium acetate, pH 4.0 to 100 μg/ml and loaded onto the Series S CM5 Sensor chip using the standard immobilization program. The final levels of immobilized YdiU were 17000-20000 response units. For the binding test for YdiU-Mn^2+^, various concentrations of Mn^2+^ were subsequently injected as analytes, and TBS (25 mM tris pH8.0, 100 mM NaCl, and 0.05% (v/v) Tween-20) was used as the running buffer. For the binding test for YdiU and nucleotide analogues, various concentrations of UMPNPP or AMPNPP were subsequently injected as analytes, and HBS (10 mM HEPES pH7.5, 150 mM NaCl_2_, 0.05% (v/v) Tween-20) was used as the running buffer. The effect of Mn^2+^ on nucleotide binding was examined by adding 50 μM Mn^2+^ to the running buffer. The binding affinity (K_D_) was calculated using the Biacore T200 evaluation software (2.0.2), and then fitting to a 1:1 binding model.

### Protein characterization with UNcle stability platform

The stabilities of YdiU before and after binding to UTP or ATP and Mn^2+^ were evaluated with an all-in-one Uncle biologics stability screening platform (Unchained Labs, Norton, MA). Purified YdiU^475^ was diluted to 2 mg/ml with a final concentration of 5 mM EDTA or 2 mM Mn^2+^ and 1 mM UTP/ATP. Then, 9 μl samples were loaded into the sample well to measure Tm with full fluorescence spectrum and size distribution with dynamic light scattering (DLS). Tm was determined using the Uncle Analysis software.

### Protein precipitation assay

A 10 μl aliquot of the indicated concentration of Mn^2+^ or/and nucleotide was added to the same volume of purified YdiU^fl^ (4 mg/ml). Samples were mixed immediately by pipetting, and then observed under microscope.

### Transmission electron microscopy

Wild type and ΔYdiU *Salmonella* were grown overnight and subcultured into fresh media. When the OD600 reached 0.4, strains were incubated for 2 min at 55°C. After recovery at 37°C for the indicated times, cells were fixed with 2.5% (w/v) glutaraldehyde at 4°C for 24 h, and untreated cells were used as the control. The samples were mixed in 1% OsO4 (w/v) at 22 °C for 2 h. After dehydration using acetone, cells were embedded using Epon 812. Specimens were then sectioned and stained with uranyl acetate and lead citrate and examined in a Hitachi transmission electron microscope with operation at 80.0 kV.

### Streptavidin pull-down assay

The streptavidin pull-down assay (SA pull-down) was performed as described in Fig.S9. Briefly, native GroEL and the GroEL Y199F, Y203F mutant were separately biotinylated by EZ-Link-Biotin and purified by desalting column. Next, 50 μg native GroEL was incubated with 20 μg YdiU^475^ (with His-tag) and 500 μM Biotin-16-UTP in a reaction buffer containing 25 mM Tris-HCl (pH7.5), 100 mM NaCl,1 mM DTT, 10 mM MnCl_2_ at 30°C for 2 h and purified by NTA Ni column and desalting column to remove free YdiU and Biotin-16-UTP. Next, 20 μg of the proteins were loaded to 50 μl High Capacity Streptavidin Agarose and washed three times with wash buffer containing 25 mM Tris-HCl (pH 8.0) and 100 mM NaCl(Rybak et al., 2004). Separately, 50 μg human ornithine carbamoyltransferase (OTC) was denatured by 8 M Urea. The denatured OTC proteins were then diluted by a factor of 50 into wash buffer and immediately added to the resin with GroEL protein and then incubated at 37 °C for 30 min. After washing twice with wash buffer, the proteins were eluted by 8 M Urea (pH 1.5) and resolved by 12% NuPAGE gel.

### Thermal resistance of wild type, ΔYdiU and YdiU-expressing *Salmonella*

The survival rates of *Salmonella strains* after heat treatment were determined as described before(Blackburn et al., 1997; Jenkins et al., 1988). First, overnight cultures were transferred into 10 ml of fresh LB medium, and when the culture OD reached 0.5, cells were harvested by centrifugation. The cell pellets were washed twice and resuspended in 0.9% NaCl solution. The samples were then divided, with some aliquots treated at 55 °C for corresponding times and the remainder serving as controls without heat treatment. Cells were plated as a dilution series on LB plates and cultured overnight at 37 °C. The next day, the viable counts of bacteria were detected as the number of colony forming units. All experiments were performed with four repeats and the results presented are mean values.

### Growth curves assay

The growth curves assay were performed using the Bioscreen C Automatic Growth Analyser (Thermo Electron Corporation) at 37℃ with a honeycomb microplate (Thermo Electron Corporation). Fresh cultured cells (OD600≈0.5) of indicated strains were diluted 100-fold in LB acidified with acetic acid to pH 6.0 and pH 5.5. 300 μL of diluted culture were added into each well and every strain was assayed in triplicates. Bacterial growth were measured at 600 nm absorbance and every 5 minutes for 24 h.

### Measurement of intracellular ATP levels

Overnight cultures of *Salmonella* were transferred into 20 ml of M9 minimal medium supplemented with 0.4% glucose, and when the culture OD reached 0.6, cells were treated at 55 °C for 1 min. Then, the cultures were divided into eight samples, which were allowed to recover at 37 °C for 5 min, 10 min, 20 min, 30 min, 45min, 60 min, 90 min, or 120 min respectively. After incubation, the OD value of each culture was detected. The cells of each sample were harvested by centrifugation at 12000 rpm for 2 min at 4°C and lysed by lysozyme. Then the ATP levels of these samples were measured by the luciferin-luciferase method(St John, 1970) following the protocol of an ATP detection kit (Beyotime, China). The average amount of intracellular ATP was obtained by calculating the ratio of total ATP and OD_600_.

## REFERENCES

Adams, P.D., Grosse-Kunstleve, R.W., Hung, L.W., Ioerger, T.R., McCoy, A.J., Moriarty, N.W., Read, R.J., Sacchettini, J.C., Sauter, N.K., and Terwilliger, T.C. (2002). PHENIX: building new software for automated crystallographic structure determination. Acta Crystallogr D Biol Crystallogr 58, 1948–1954.

Adler, S.P., Purich, D., and Stadtman, E.R. (1975). Cascade control of Escherichia coli glutamine synthetase. Properties of the PII regulatory protein and the uridylyltransferase-uridylyl-removing enzyme. The Journal of biological chemistry 250, 6264–6272.

Balleza, E., Lopez-Bojorquez, L.N., Martinez-Antonio, A., Resendis-Antonio, O., Lozada-Chavez, I., Balderas-Martinez, Y.I., Encarnacion, S., and Collado-Vides, J. (2009). Regulation by transcription factors in bacteria: beyond description. FEMS microbiology reviews 33, 133–151.

Blackburn, C.W., Curtis, L.M., Humpheson, L., Billon, C., and McClure, P.J. (1997). Development of thermal inactivation models for Salmonella enteritidis and Escherichia coli O157:H7 with temperature, pH and NaCl as controlling factors. International journal of food microbiology 38, 31–44.

Browning, D.F., and Busby, S.J. (2004). The regulation of bacterial transcription initiation. Nature reviews. Microbiology 2, 57–65.

Buckle, A.M., Zahn, R., and Fersht, A.R. (1997). A structural model for GroEL-polypeptide recognition. Proceedings of the National Academy of Sciences of the United States of America 94, 3571–3575.

Cain, J.A., Solis, N., and Cordwell, S.J. (2014). Beyond gene expression: the impact of protein post-translational modifications in bacteria. Journal of proteomics 97, 265–286.

Carabetta, V.J., and Cristea, I.M. (2017). Regulation, Function, and Detection of Protein Acetylation in Bacteria. Journal of bacteriology 199.

Case, D.A., Cheatham, T.E., 3rd, Darden, T., Gohlke, H., Luo, R., Merz, K.M., Jr., Onufriev, A., Simmerling, C., Wang, B., and Woods, R.J. (2005). The Amber biomolecular simulation programs. Journal of computational chemistry 26, 1668–1688.

Casey, A.K., and Orth, K. (2018). Enzymes Involved in AMPylation and deAMPylation. Chem Rev 118, 1199–1215.

Coleman, T.G., Mesick, H.C., and Darby, R.L. (1977). Numerical integration: a method for improving solution stability in models of the circulation. Annals of biomedical engineering 5, 322–328.

Datsenko, K.A., and Wanner, B.L. (2000). One-step inactivation of chromosomal genes in Escherichia coli K-12 using PCR products. Proceedings of the National Academy of Sciences of the United States of America 97, 6640–6645.

Diamant, S., and Goloubinoff, P. (1998). Temperature-controlled activity of DnaK-DnaJ-GrpE chaperones: protein-folding arrest and recovery during and after heat shock depends on the substrate protein and the GrpE concentration. Biochemistry 37, 9688–9694.

Dudkiewicz, M., Szczepinska, T., Grynberg, M., and Pawlowski, K. (2012). A novel protein kinase-like domain in a selenoprotein, widespread in the tree of life. PloS one 7, e32138.

Dukan, S., and Nystrom, T. (1998). Bacterial senescence: stasis results in increased and differential oxidation of cytoplasmic proteins leading to developmental induction of the heat shock regulon. Genes & development 12, 3431–3441.

Emsley, P., and Cowtan, K. (2004). Coot: model-building tools for molecular graphics. Acta Crystallogr D Biol Crystallogr 60, 2126–2132.

Erasmus, D.J., van der Merwe, G.K., and van Vuuren, H.J. (2003). Genome-wide expression analyses: Metabolic adaptation of Saccharomyces cerevisiae to high sugar stress. FEMS yeast research 3, 375–399.

Feng, F., Yang, F., Rong, W., Wu, X., Zhang, J., Chen, S., He, C., and Zhou, J.M. (2012). A Xanthomonas uridine 5’-monophosphate transferase inhibits plant immune kinases. Nature 485, 114–118.

Fenton, W.A., Kashi, Y., Furtak, K., and Horwich, A.L. (1994). Residues in chaperonin GroEL required for polypeptide binding and release. Nature 371, 614–619.

Galperin, M.Y., and Koonin, E.V. (2004). ’Conserved hypothetical’ proteins: prioritization of targets for experimental study. Nucleic acids research 32, 5452–5463.

Galperin, M.Y., and Koonin, E.V. (2010). From complete genome sequence to ‘complete’ understanding? Trends Biotechnol 28, 398–406.

Goodsell, D.S., Morris, G.M., and Olson, A.J. (1996). Automated docking of flexible ligands: applications of AutoDock. Journal of molecular recognition : JMR 9, 1–5.

Gragerov, A., Nudler, E., Komissarova, N., Gaitanaris, G.A., Gottesman, M.E., and Nikiforov, V. (1992). Cooperation of GroEL/GroES and DnaK/DnaJ heat shock proteins in preventing protein misfolding in Escherichia coli. Proceedings of the National Academy of Sciences of the United States of America 89, 10341–10344.

Grangeasse, C., Stulke, J., and Mijakovic, I. (2015). Regulatory potential of post-translational modifications in bacteria. Frontiers in microbiology 6, 500.

Ham, H., Woolery, A.R., Tracy, C., Stenesen, D., Kramer, H., and Orth, K. (2014). Unfolded protein response-regulated Drosophila Fic (dFic) protein reversibly AMPylates BiP chaperone during endoplasmic reticulum homeostasis. The Journal of biological chemistry 289, 36059–36069.

Harrison, C.J., Hayer-Hartl, M., Di Liberto, M., Hartl, F., and Kuriyan, J. (1997). Crystal structure of the nucleotide exchange factor GrpE bound to the ATPase domain of the molecular chaperone DnaK. Science 276, 431–435.

Helmann, J.D., and Chamberlin, M.J. (1988). Structure and function of bacterial sigma factors. Annual review of biochemistry 57, 839–872.

Hendrickson, W.A., Horton, J.R., and LeMaster, D.M. (1990). Selenomethionyl proteins produced for analysis by multiwavelength anomalous diffraction (MAD): a vehicle for direct determination of three-dimensional structure. EMBO J 9, 1665–1672.

Hoch, J.A. (2000). Two-component and phosphorelay signal transduction. Current opinion in microbiology 3, 165–170.

Hsu-Ming, W., Naito, K., Kinoshita, Y., Kobayashi, H., Honjoh, K., Tashiro, K., and Miyamoto, S. (2012). Changes in transcription during recovery from heat injury in Salmonella typhimurium and effects of BCAA on recovery. Amino acids 42, 2059–2066.

Hu, L.I., Lima, B.P., and Wolfe, A.J. (2010). Bacterial protein acetylation: the dawning of a new age. Molecular microbiology 77, 15–21.

Humphrey, W., Dalke, A., and Schulten, K. (1996). VMD: visual molecular dynamics. Journal of molecular graphics 14, 33-38, 27–38.

Jakubovics, N.S., and Jenkinson, H.F. (2001). Out of the iron age: new insights into the critical role of manganese homeostasis in bacteria. Microbiology 147, 1709–1718.

Jenkins, D.E., Schultz, J.E., and Matin, A. (1988). Starvation-induced cross protection against heat or H2O2 challenge in Escherichia coli. Journal of bacteriology 170, 3910–3914.

Jones, J.D., and O’Connor, C.D. (2011). Protein acetylation in prokaryotes. Proteomics 11, 3012–3022.

Kehres, D.G., Zaharik, M.L., Finlay, B.B., and Maguire, M.E. (2000). The NRAMP proteins of Salmonella typhimurium and Escherichia coli are selective manganese transporters involved in the response to reactive oxygen. Molecular microbiology 36, 1085–1100.

Lee, S., Sowa, M.E., Watanabe, Y.H., Sigler, P.B., Chiu, W., Yoshida, M., and Tsai, F.T. (2003). The structure of ClpB: a molecular chaperone that rescues proteins from an aggregated state. Cell 115, 229–240.

Li, B., Yue, Y., Yuan, Z., Zhang, F., Li, P., Song, N., Lin, W., Liu, Y., Yang, Y., Li, Z., et al. (2017). Salmonella STM1697 coordinates flagella biogenesis and virulence by restricting flagellar master protein FlhD4C2 from recruiting RNA polymerase. Nucleic acids research 45, 9976–9989.

Li, P., and Merz, K.M., Jr. (2016). MCPB.py: A Python Based Metal Center Parameter Builder. Journal of chemical information and modeling 56, 599–604.

Liberek, K., Marszalek, J., Ang, D., Georgopoulos, C., and Zylicz, M. (1991). Escherichia coli DnaJ and GrpE heat shock proteins jointly stimulate ATPase activity of DnaK. Proceedings of the National Academy of Sciences of the United States of America 88, 2874–2878.

Maier, J.A., Martinez, C., Kasavajhala, K., Wickstrom, L., Hauser, K.E., and Simmerling, C. (2015). ff14SB: Improving the Accuracy of Protein Side Chain and Backbone Parameters from ff99SB. Journal of chemical theory and computation 11, 3696–3713.

McCarty, J.S., and Walker, G.C. (1991). DnaK as a thermometer: threonine-199 is site of autophosphorylation and is critical for ATPase activity. Proceedings of the National Academy of Sciences of the United States of America 88, 9513–9517.

Otwinowski, Z., and Minor, W. (1997). Processing of X-ray diffraction data collected in oscillation mode. Methods Enzymol 276, 307–326.

Phadtare, S., and Inouye, M. (2004). Genome-wide transcriptional analysis of the cold shock response in wild-type and cold-sensitive, quadruple-csp-deletion strains of Escherichia coli. Journal of bacteriology 186, 7007–7014.

Price, D.J., and Brooks, C.L., 3rd (2004). A modified TIP3P water potential for simulation with Ewald summation. The Journal of chemical physics 121, 10096–10103.

Roux, K.J., Kim, D.I., and Burke, B. (2013). BioID: a screen for protein-protein interactions. Curr Protoc Protein Sci 74, Unit 19 23.

Rybak, J.N., Scheurer, S.B., Neri, D., and Elia, G. (2004). Purification of biotinylated proteins on streptavidin resin: a protocol for quantitative elution. Proteomics 4, 2296–2299.

Sanyal, A., Chen, A.J., Nakayasu, E.S., Lazar, C.S., Zbornik, E.A., Worby, C.A., Koller, A., and Mattoo, S. (2015). A novel link between Fic (filamentation induced by cAMP)-mediated adenylylation/AMPylation and the unfolded protein response. The Journal of biological chemistry 290, 8482–8499.

Schwarz, F., and Aebi, M. (2011). Mechanisms and principles of N-linked protein glycosylation. Current opinion in structural biology 21, 576–582.

Sharma, S.K., De los Rios, P., Christen, P., Lustig, A., and Goloubinoff, P. (2010). The kinetic parameters and energy cost of the Hsp70 chaperone as a polypeptide unfoldase. Nature chemical biology 6, 914–920.

Sherman, M., and Goldberg, A.L. (1994). Heat shock-induced phosphorylation of GroEL alters its binding and dissociation from unfolded proteins. The Journal of biological chemistry 269, 31479–31483.

Shiau, A.K., Harris, S.F., Southworth, D.R., and Agard, D.A. (2006). Structural Analysis of E. coli hsp90 reveals dramatic nucleotide-dependent conformational rearrangements. Cell 127, 329–340.

Sreelatha, A., Yee, S.S., Lopez, V.A., Park, B.C., Kinch, L.N., Pilch, S., Servage, K.A., Zhang, J., Jiou, J., Karasiewicz-Urbanska, M., et al. (2018). Protein AMPylation by an Evolutionarily Conserved Pseudokinase. Cell 175, 809–821 e819.

St John, J.B. (1970). Determination of ATP in Chlorella with the luciferin-luciferase enzyme system. Analytical biochemistry 37, 409–416.

Stock, A.M., Robinson, V.L., and Goudreau, P.N. (2000). Two-component signal transduction. Annual review of biochemistry 69, 183–215.

Terwilliger, T.C. (2002). Automated structure solution, density modification and model building. Acta Crystallogr D Biol Crystallogr 58, 1937–1940.

Thomas, J.G., and Baneyx, F. (2000). ClpB and HtpG facilitate de novo protein folding in stressed Escherichia coli cells. Molecular microbiology 36, 1360–1370.

Truttmann, M.C., Pincus, D., and Ploegh, H.L. (2018). Chaperone AMPylation modulates aggregation and toxicity of neurodegenerative disease-associated polypeptides. Proceedings of the National Academy of Sciences of the United States of America 115, E5008–E5017.

Truttmann, M.C., Zheng, X., Hanke, L., Damon, J.R., Grootveld, M., Krakowiak, J., Pincus, D., and Ploegh, H.L. (2017). Unrestrained AMPylation targets cytosolic chaperones and activates the heat shock response. Proceedings of the National Academy of Sciences of the United States of America 114, E152–E160.

Wada, T., Morizane, T., Abo, T., Tominaga, A., Inoue-Tanaka, K., and Kutsukake, K. (2011). EAL domain protein YdiV acts as an anti-FlhD4C2 factor responsible for nutritional control of the flagellar regulon in Salmonella enterica Serovar Typhimurium. Journal of bacteriology 193, 1600–1611.

Worby, C.A., Mattoo, S., Kruger, R.P., Corbeil, L.B., Koller, A., Mendez, J.C., Zekarias, B., Lazar, C., and Dixon, J.E. (2009). The fic domain: regulation of cell signaling by adenylylation. Mol Cell 34, 93–103.

Yarbrough, M.L., Li, Y., Kinch, L.N., Grishin, N.V., Ball, H.L., and Orth, K. (2009). AMPylation of Rho GTPases by Vibrio VopS disrupts effector binding and downstream signaling. Science 323, 269–272.

Zhang, H., Yang, J., Wu, S., Gong, W., Chen, C., and Perrett, S. (2016). Glutathionylation of the Bacterial Hsp70 Chaperone DnaK Provides a Link between Oxidative Stress and the Heat Shock Response. The Journal of biological chemistry 291, 6967–6981.

