## supplemental informations for "The YdiU Domain Modulates Bacterial Stress Signaling through Mn^2+^-dependent UMPylation"

### Supplementary Fig.S1-14

### Supplementary Table S1-6

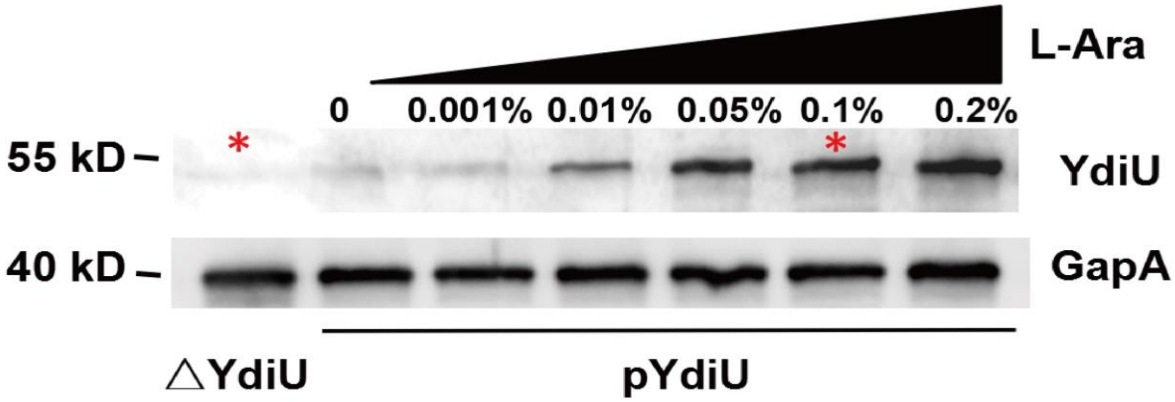


**Fig.S1. The expression of YdiU in pYdiU strain inducing with 0-0.2% L-arabinose.**

The expression of YdiU in LB medium with indicated concentration of L-arabinose was quantified by western blot. ΔydiU strain (Lane1) was used as a negative control. GapA was used as a loading control. The strains and conditions used for subsequent MS were highlighted in red asterisks.

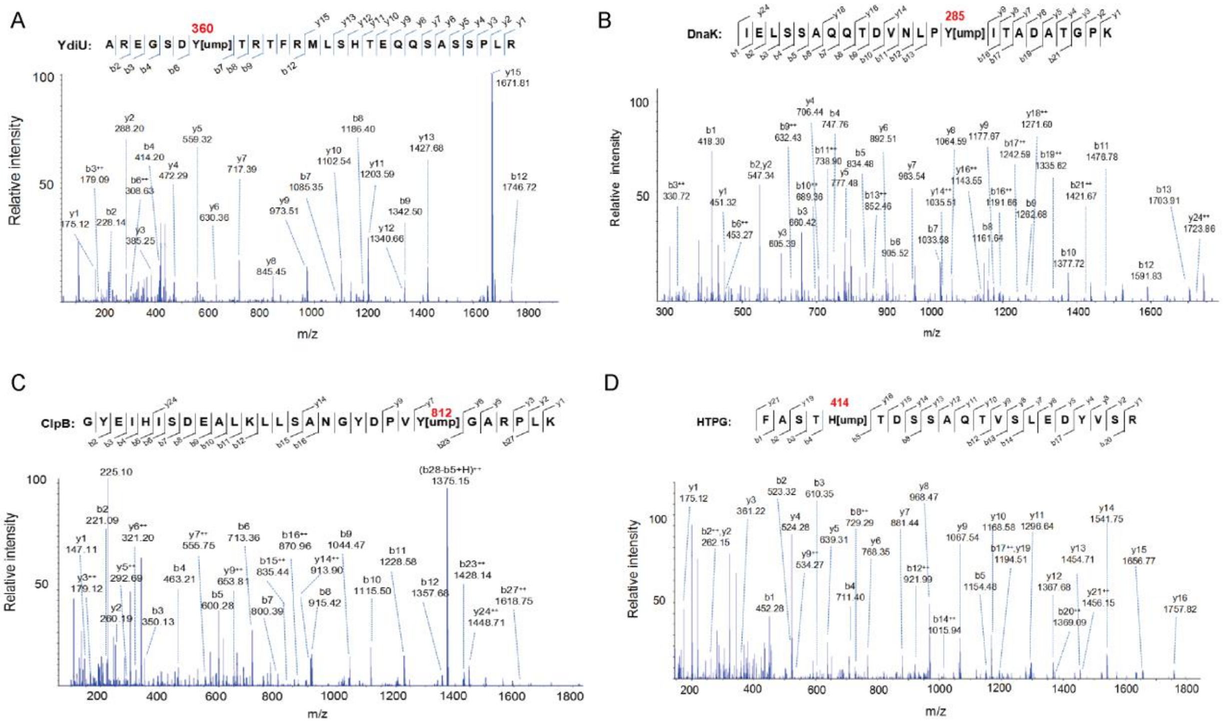


**Fig.S2. Electrospray ionization MS/MS spectra of UMPylated peptides identified in YdiU-expressing *Salmonella*, Related to Fig.2.**

(A) YdiU peptide with an UMPylated Tyr360. The increased mass of 306.025 daltons was detected after b7, unambiguously confirming Tyr360 was the UMPylated residue. (B) DnaK peptide with an UMPylated Tyr285. (C) ClpB peptide with an UMPylated Tyr812. (D) HtpG peptide with an UMPylated His414.

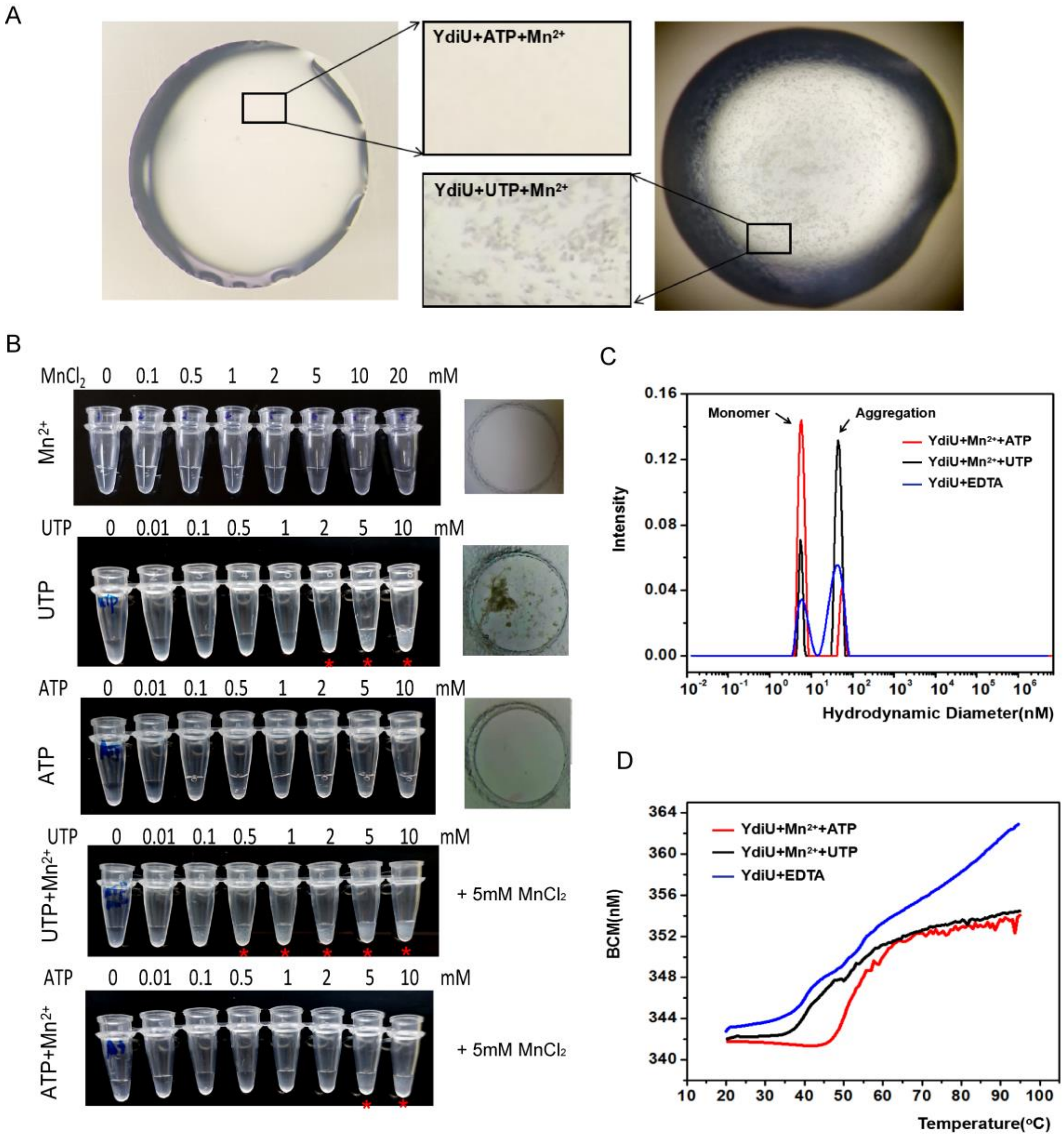


**Fig.S3. UTP and $Mn^{2+}$-binding turns YdiU to an aggregation-prone state.**

(A) Precipitate was observed in the YdiU-UTP-$Mn^{2+}$ but not in the YdiU-ATP-$Mn^{2+}$ in the same condition. This picture is a representative of multiple similar drops in different conditions. (B) Protein precipitation assay performed as described in Methods. The last samples in each row were also observed under microscope. (C) Hydrodynamic diameter distributions of YdiU-ATP-$Mn^{2+}$ and YdiU-UTP-$Mn^{2+}$ on DLS. (D) $T_m$ of YdiU in apo state or nucleic acid bound state obtained from DSF with the UNcle platform.

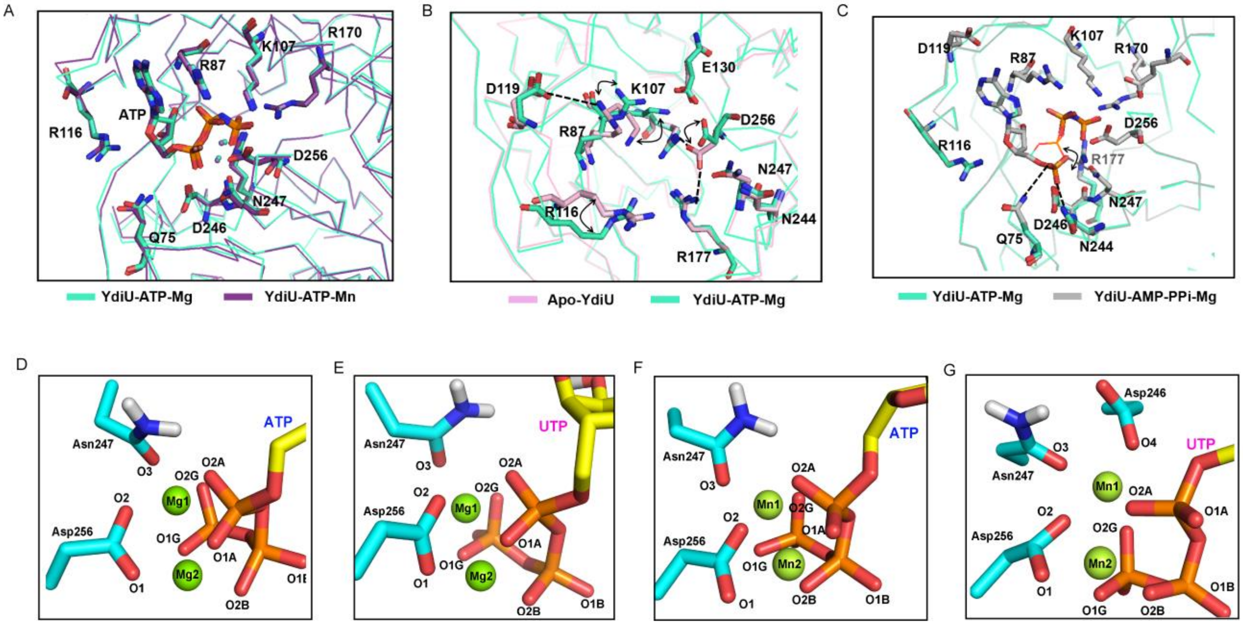


**Fig.S4. Structural comparison reveals the essential residues involved in nucleotide and ion-binding .**

(A) Superposition of YdiU-ATP-$Mg^{2+}$ and YdiU-ATP-$Mn^{2+}$. No dramatically conformation change can be observed. (B) Superposition of Apo-YdiU and YdiU-ATP. The residues in the active center are shown in sticks in indicated color. The conformation shifts were marked by double-heated black arrows. (C) Superposition of YdiU-ATP and YdiU-AMP-PPi. The residues in the active center are shown in sticks in indicated color. The conformation shift in α-phospate group was highlighted by double-heated black arrows. The interactions between phospate of AMP and Q75, N244 and N247 were indicated as black dashed lines. (D-G) The nucleotide and ion binding models generated by molecular dynamics simulations. Remarkably, Asp246 provides an extra bond interaction with $Mn^{2+}$ in the model of YdiU-UTP-$Mn^{2+}$.

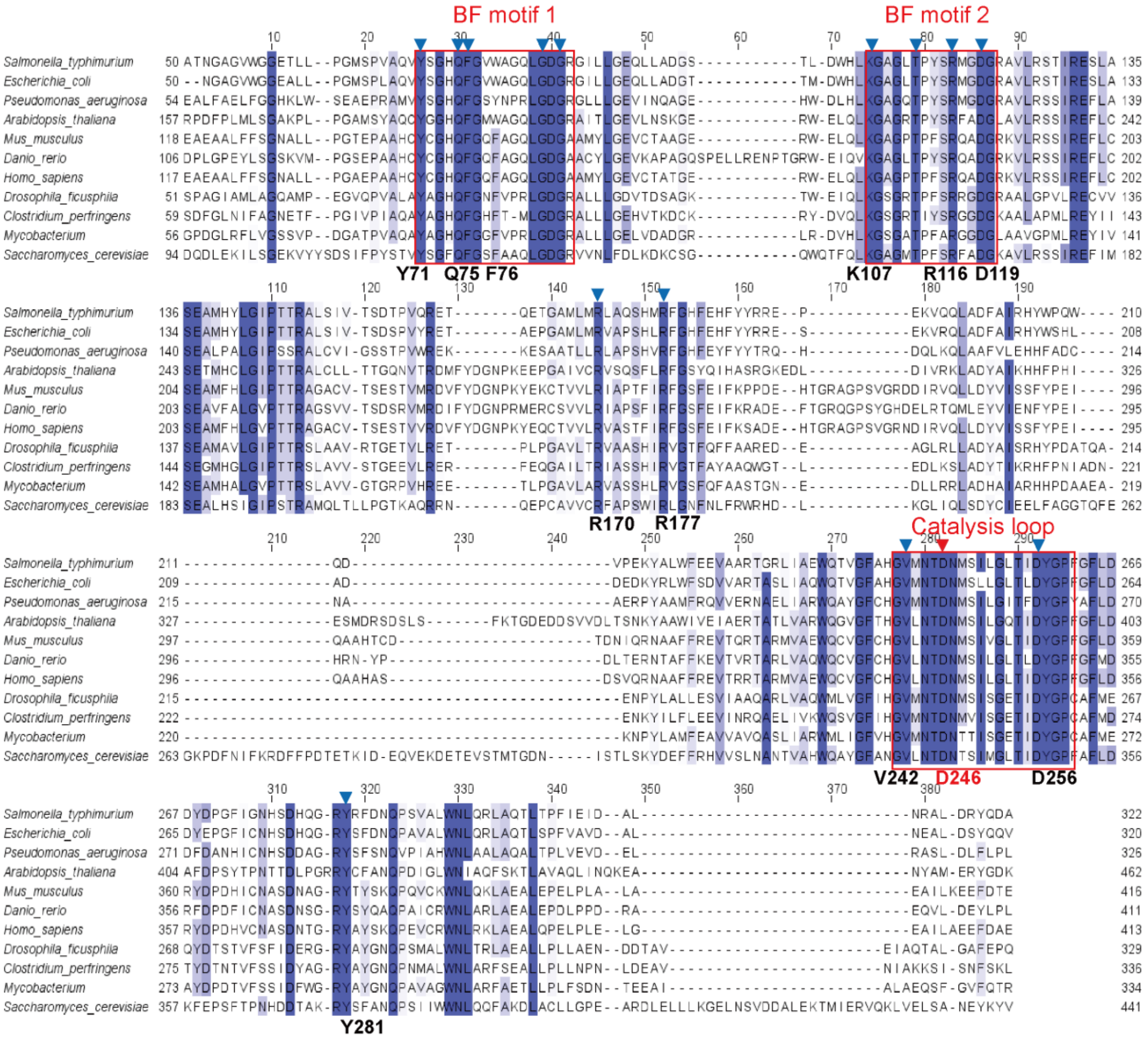


**Fig.S5. Sequence alignment of YdiU homologues.**

The YdiU family was identified by NCBI conserved domains, as aligned by T-coffee and Clustal W, and colored by JalView. The key residues identified in this study are labeled. The conserved basic fixed motifs and catalysis loop were highlighted by red box.

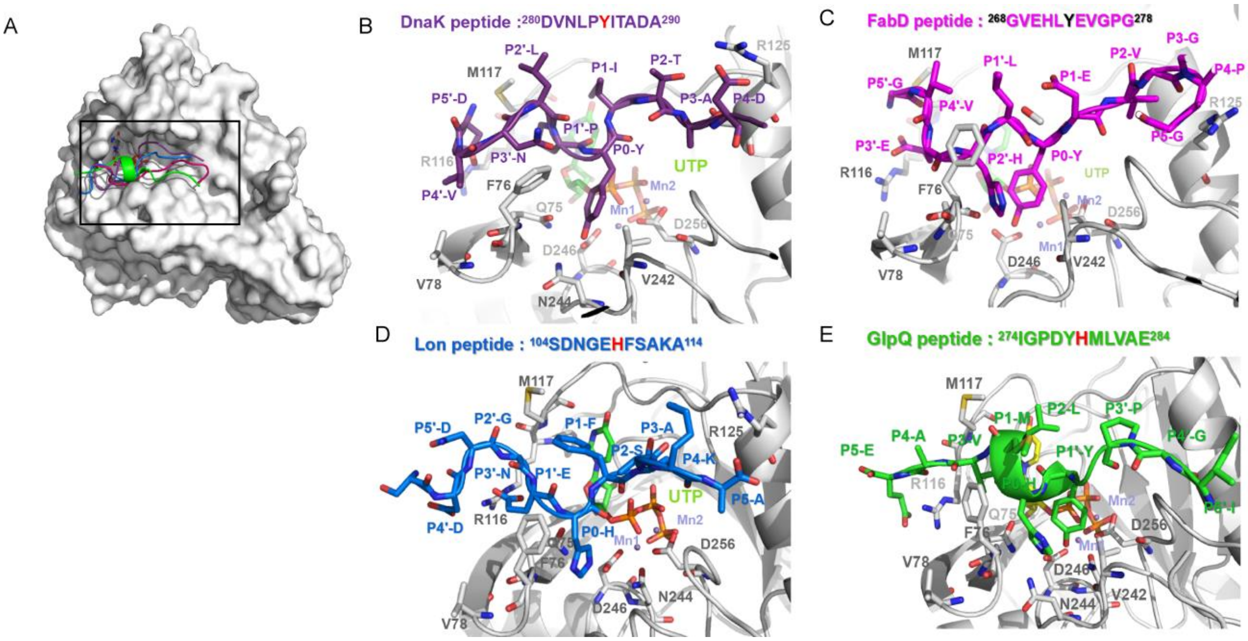


**Fig.S6. Models of YdiU-UTP-Mn-peptide complex, related to Fig.4L.**

(A-E) Four UMPylated peptides identified in YdiU-expressing *Salmonella* were docked into the structure of YdiU-UTP-$Mn^{2+}$. (A) The overall view of substrate binding site. YdiU is shown in surface mode and four peptides are shown in cartoon mode. (B-E) The zoomed view of YdiU-UTP-Mn-peptides. Peptides were shown in stick with corresponding color. The overall structures of YdiU were shown in cartoon, and the residues involved in peptides-binding were shown in stick. The identified UMPylated site was highlighted in the peptide sequence above the structure chart.

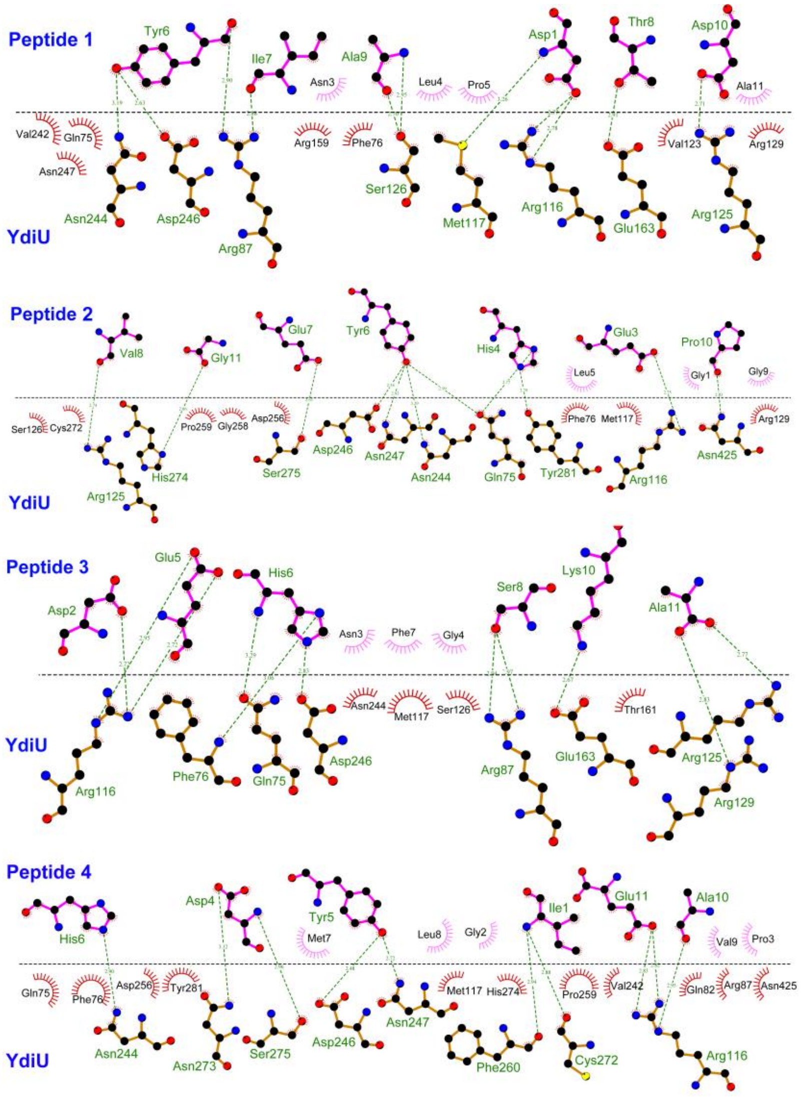


**Fig.S7. Interaction surface of YdiU and peptides, related to Fig.4L and Fig.S6.**

Four UMPylated peptides identified in YdiU-expressing *Salmonella* were docked into the structure of YdiU-UTP-$Mn^{2+}$. The interactions between YdiU and peptides were analyzed using Ligplus. The hydrogen bond interactions are labeled by green dotted line. Peptide1: DVNLPYITADA from DnaK; Peptide2: GVEHLYEVGPG from FabD; Peptide3: SDNGEHFSAKA from Lon; Peptide4: IGPDYHMLVAE from GlpQ.

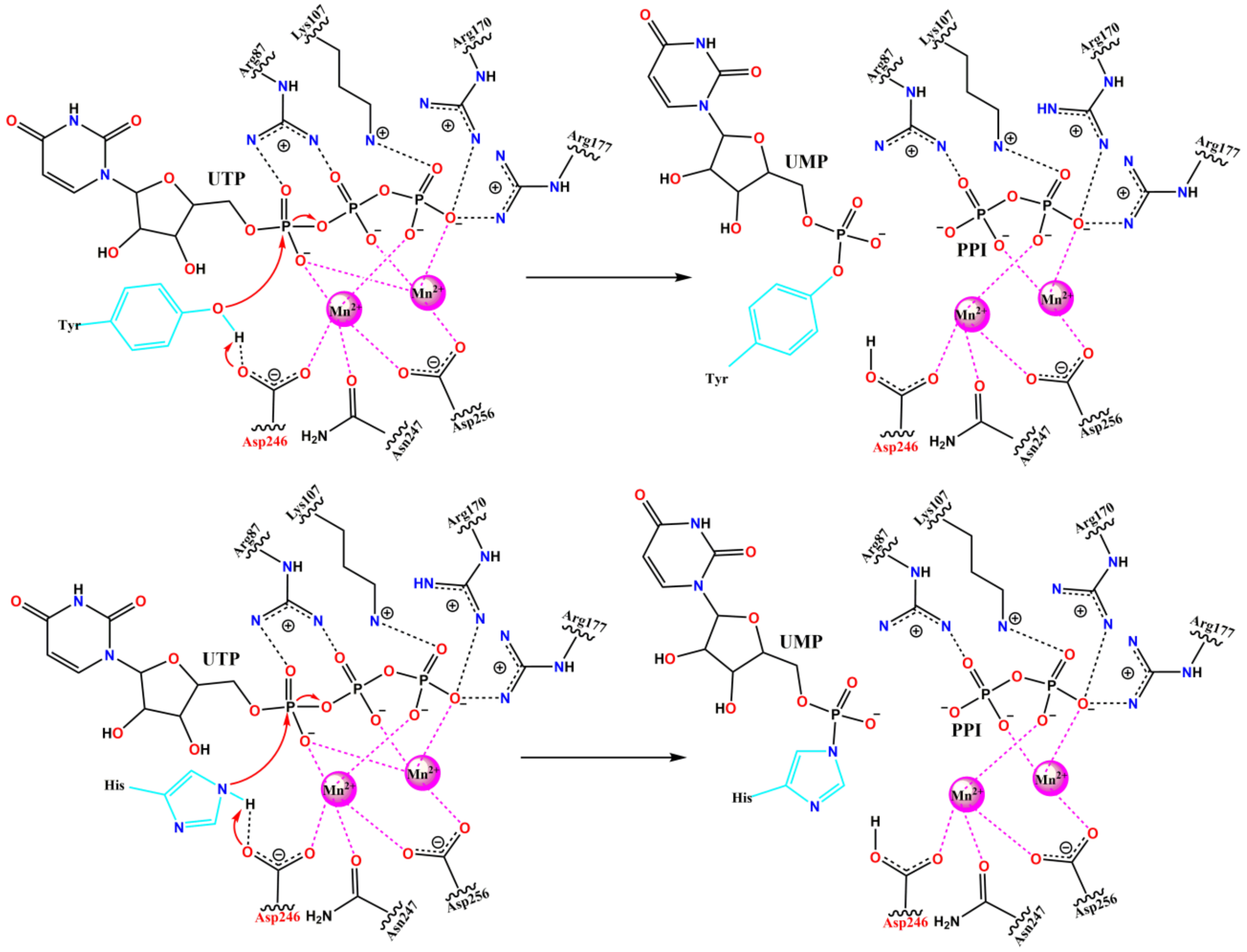


**Fig.S8. Catalytic mechanism of YdiU-mediated UMPylation, related to Fig.4M.**

The proposed mechanism of YdiU-mediated UMPylation. Asp 246 acts as the general base and activates the oxygen of the hydroxyl group from Tyr or the nitrogen from His for nucleophilic attack, then the UMP moiety of UTP transfers to the Tyr or His residue of substrate.

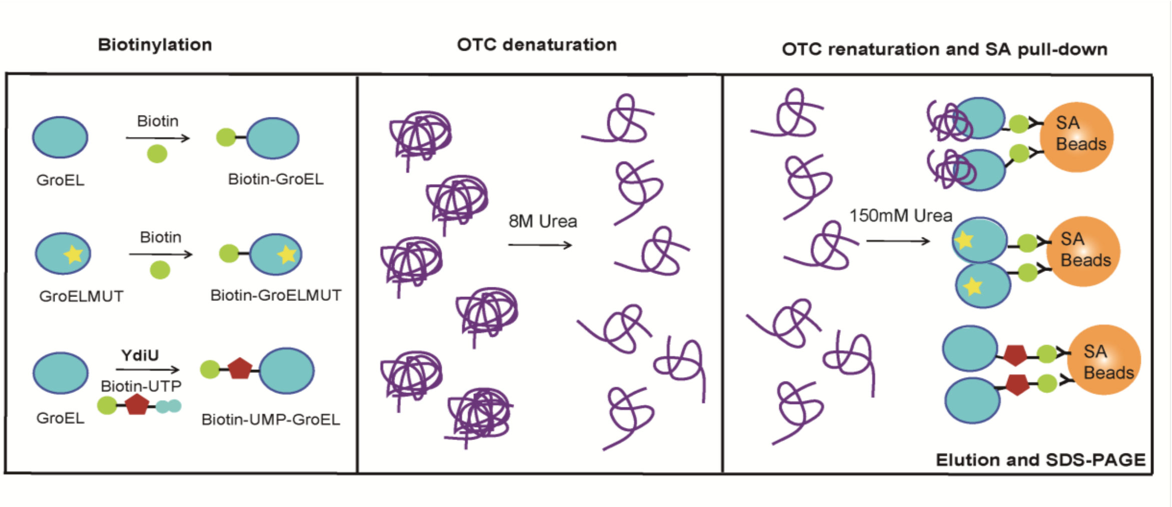


**Fig.S9. Schematic of the streptavidin pull-down approach, related to Fig.6C.**

GroEL were biotinylated by biotin-linker or by biotin-16-UTP and YdiU, GroEL mutant was biotinylated by biotin-linker. The biotinylated proteins were bound to Streptavidin Agarose. OTC was denatured by 8 M Urea. The production was diluted by a factor of 50 and added to the resin with GroEL. After wash for three times, the elution was resolved by SDS-PAGE.

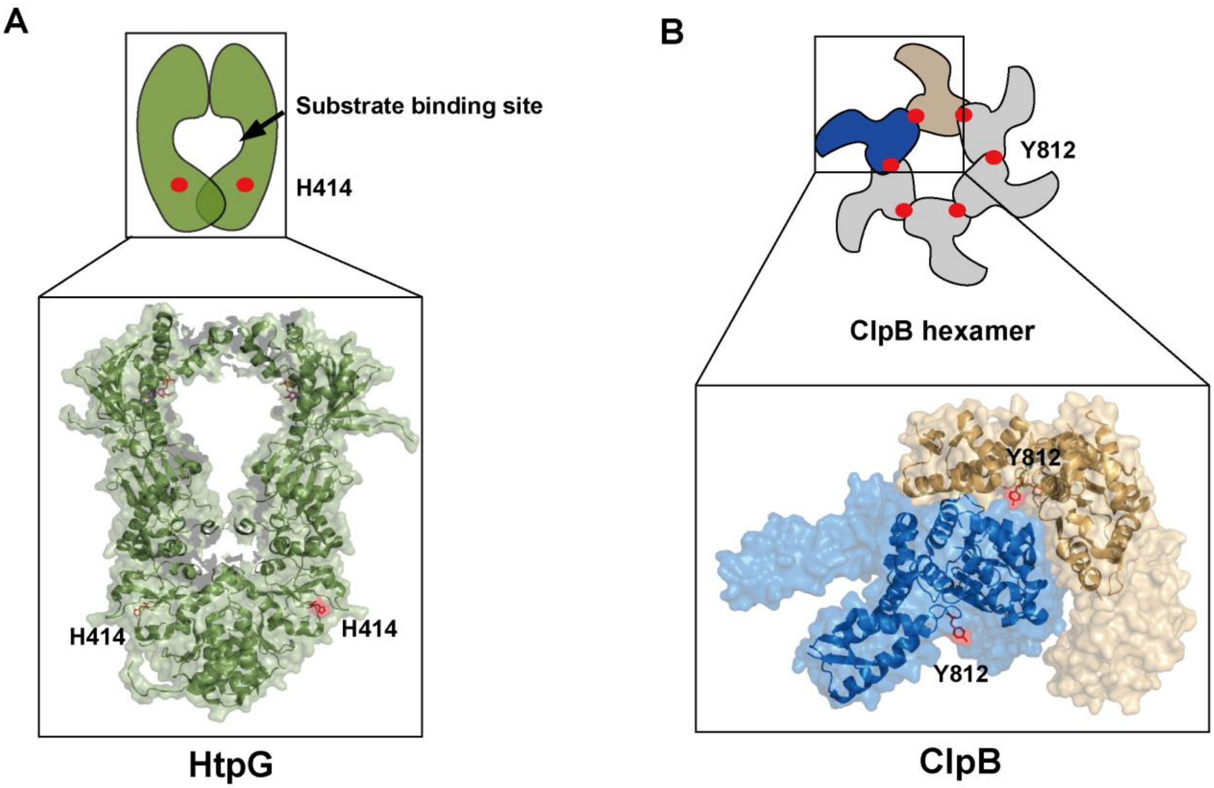


**Fig.S10. The structures of HtpG and ClpB indicating their UMPylation sites.**

(A) Schematic representation of HtpG dimer. The structure of HtpG is shown in cartoon and surface mode. H414, the UMPylated site of HtpG, is highlighted in red stick. PDB code:2IOP.

(B) Schematic representation of ClpB hexamer. The structure of two subunits of ClpB hexamer is shown in cartoon and surface mode. Y812, the UMPylated site of ClpB identified *in vivo,* is highlighted in red stick. PDB code:4CIU.

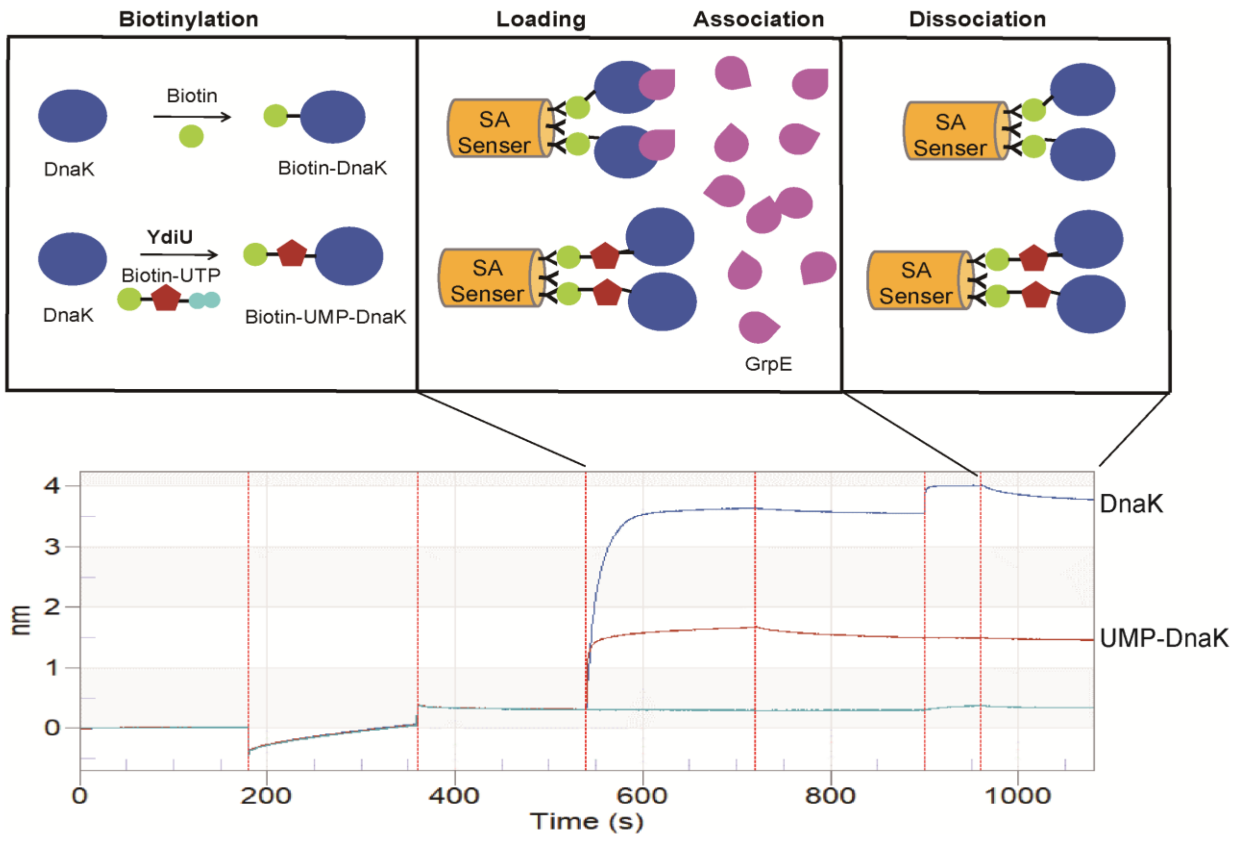


**Fig.S11. Schematic of the streptavidin-based BLI approach, related to Fig.6E.**

DnaK proteins were biotinylated either by biotin-linker or by biotin-16-UTP and YdiU, and then separately loaded to streptavidin sensors. The sensors were introduced to GrpE to measure the association response and changed to PBS buffer to measure dissociation. The figure below presents complete curve of this experiment.

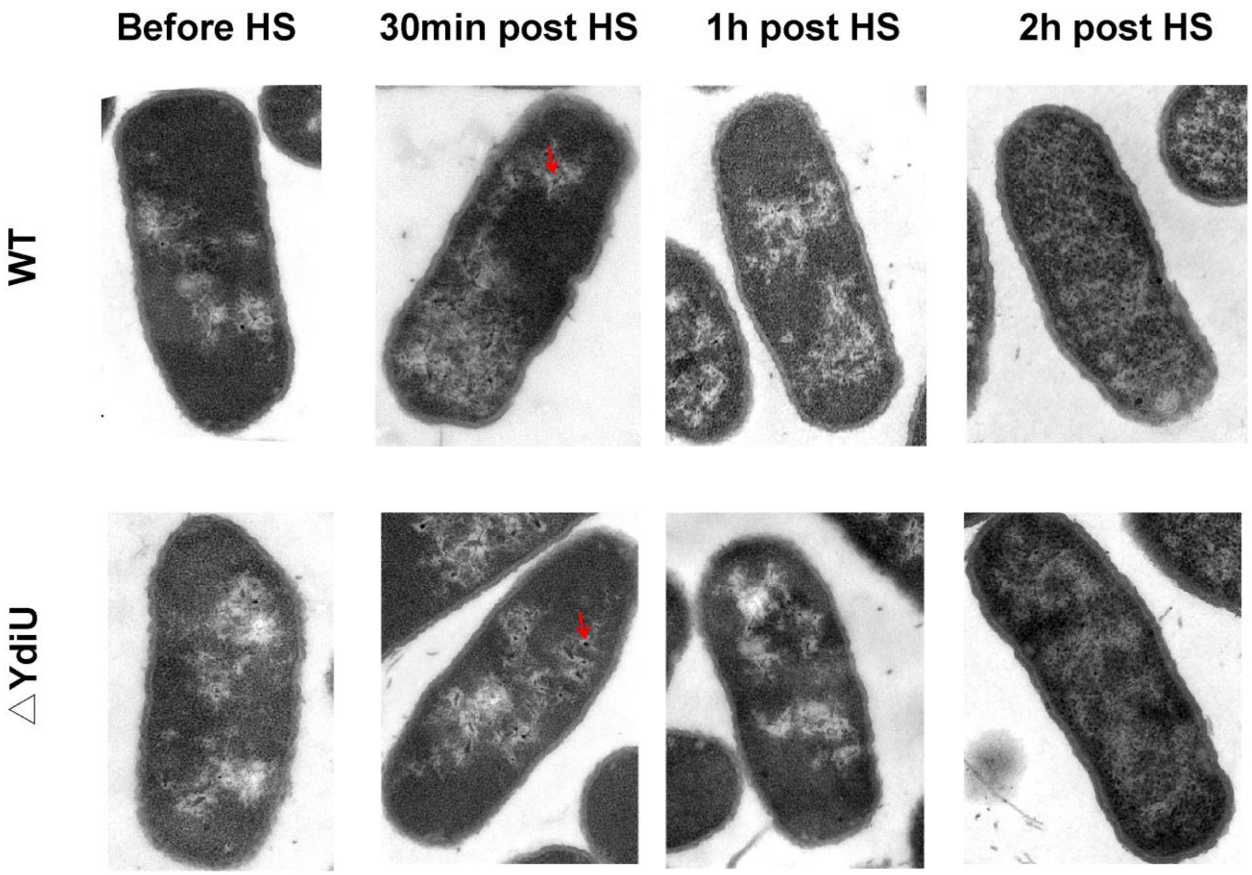


**Fig.S12. Electron micrographs of wild-type and ΔYdiU *Salmonella* following by heat shock.** Wild type and ΔYdiU *Salmonella* were treated for 2 min at 55°C, then the cells following recovery at 37°C for the indicated time were observed using transmission electron microscopy. Inclusion bodies are indicated by red arrowheads.

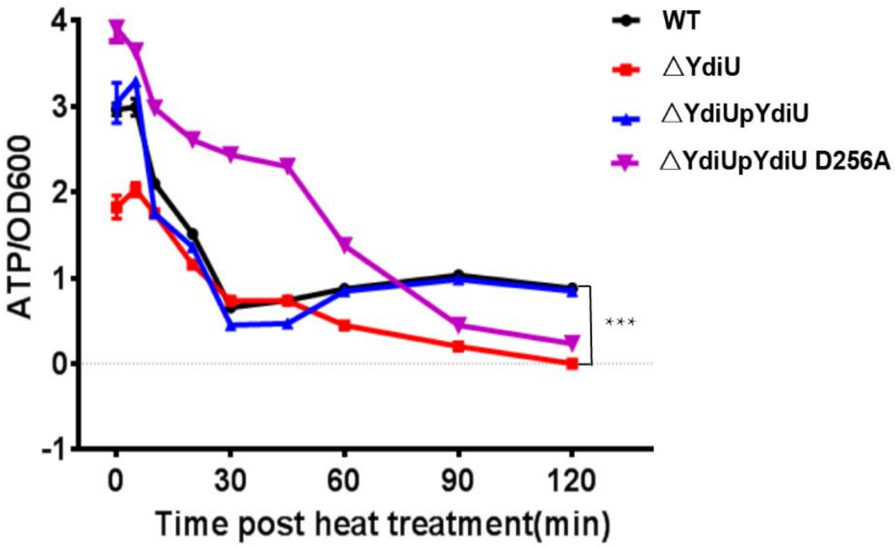


**Fig.S13. ATP levels of *Salmonella* strains during recovery from heat injury, related to Fig.7C.**

ATP levels of the indicated strains of *Salmonella* during recovery at the indicated time after heat injury. The average amount of intracellular ATP was obtained by calculating the ratio of total ATP and OD600. All experiments were performed as three repeats and the mean values and error bar are presented. Statistical significance is indicated by ***P < 0.001 using a t-test.

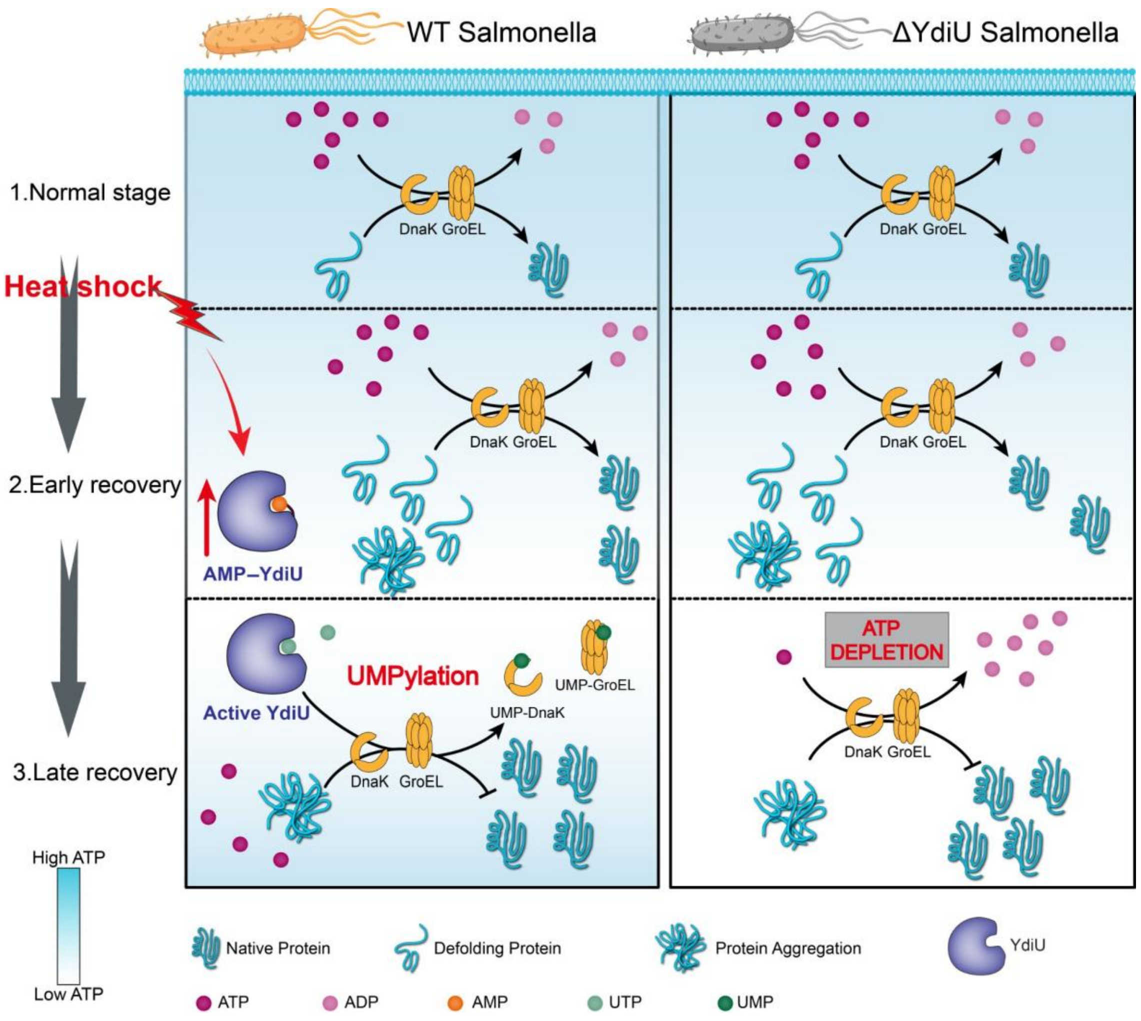


**Fig.S14. The model of YdiU-mediated chaperone-switch during heat stress.**

Heat produces misfolding proteins in cells. Firstly, Chaperones are expressed, they repair the misfolded proteins using ATP. Subsequently, YdiU is expressed. When the intracellular concentration of ATP is in a high level, YdiU is self-AMPylated and inactive. In the later stage of recovery, most of misfolded proteins were refolded by chaperones, consuming lots of ATP. Under ATP-limited condition, YdiU is activated as an UMPylator to UMPylate and inactive chaperones, thereby preventing stress-induced ATP depletion. The YdiU knockout strain will die of ATP depletion causing by excessive ATP consumption of chaperones.

### Table S1 UMPylated peptides identified in YdiU-expressing *Salmonella*

| | Pro | Sequence | Modifications | Prec MW | Theor MW | dMass | Theor z |
|---|---|---|---|---|---|---|---|
| 1 | DNAK | IELSSAQQTDVNLPYITADATGPK | PhosphoUridine(Y)@15 | 3445.898 | 3445.711 | 0.185455 | 3 |
| 2 | SYI | YGLETANPVGPDGTYLPGTYPTLDGVNVFK | PhosphoUridine(Y)@15 | 4069.157 | 4068.986 | 0.170564 | 3 |
| 3 | A0A0F6B416 | QVAEYADGIGPDYHMLVAEGSTK | PhosphoUridine(H)@14 | 3364.754 | 3364.578 | 0.175942 | 3 |
| 4 | LEPA | EYDLDLITTAPTVVYEVETTAK | PhosphoUridine(Y)@15 | 3384.857 | 3384.673 | 0.184984 | 3 |
| 5 | OMPC | YDANNIYLAAQYSQTYNATR | PhosphoUridine(Y)@7 | 2950.494 | 2950.297 | 0.19781 | 3 |
| 6 | OMPC | YDANNIYLAAQYSQTYNATR | PhosphoUridine(Y)@7 | 2949.517 | 2949.313 | 0.204678 | 3 |
| 7 | METK | IVGIDAVVLSTQHAEDIDQK | PhosphoUridine(H)@13 | 3064.741 | 3064.558 | 0.183467 | 2 |
| 8 | G3P | VINDNFGIIEGLMTTVHATTATQK | PhosphoUridine(H)@17 | 3488.897 | 3488.736 | 0.16218 | 3 |
| 9 | ATPB | MPSAVGYQPTLAEEMGVLQER | PhosphoUridine(Y)@7 | 2915.522 | 2915.339 | 0.183559 | 3 |
| 10 | PURA | LLLSEACPLILDYHVALDNAR | PhosphoUridine(Y)@13 | 3006.665 | 3006.472 | 0.193836 | 3 |
| 11 | PURA | LLLSEACPLILDYHVALDNAR | PhosphoUridine(Y)@13 | 3005.681 | 3005.488 | 0.194774 | 3 |
| 12 | ATPG | YVESQVYQGVVENLASEQAAR | PhosphoUridine(Y)@7 | 2949.557 | 2949.37 | 0.186534 | 3 |
| 13 | A0A0F6B033 | DGFVLGDGAGMLVLEEYEHAK | PhosphoUridine(Y)@17 | 3163.688 | 3163.503 | 0.183423 | 3 |
| 14 | RPOB | INPIEDMPYDENGTPVDIVLNPLGVPSR | PhosphoUridine(Y)@9 | 3689.918 | 3689.748 | 0.169938 | 3 |
| 15 | RPOB | QNQLEQLAEQYDELKHEFEK | PhosphoUridine(Y)@11 | 3737.033 | 3736.839 | 0.193681 | 3 |
| 16 | A0A0F6AXW7 | FASTHTDSSAQTVSLEDYVSR | PhosphoUridine(H)@5 | 2910.484 | 2910.286 | 0.19768 | 3 |
| 17 | G3P | VINDNFGIIEGLMTTVHATTATQK | PhosphoUridine(H)@17 | 3488.906 | 3488.736 | 0.169157 | 3 |
| 18 | PNP | EGDNYVVLSDILGDEDHLGDMDFK | PhosphoUridine(H)@17 | 3609.815 | 3609.632 | 0.184441 | 3 |
| 19 | GLMS | GLDASIEHDIVHGLQALPSR | PhosphoUridine(H)@8 | 2737.522 | 2737.338 | 0.183957 | 3 |
| 20 | METK | IVGIDAVVLSTQHAEDIDQK | PhosphoUridine(H)@13 | 3064.745 | 3064.558 | 0.187833 | 3 |
| 21 | ARAB | NPSLDHLPVVLDWFNGR | PhosphoUridine(H)@6 | 2589.405 | 2589.221 | 0.184661 | 3 |
| 22 | MURA | AEIESNTVICHGVETLSGAQVMATDLR | PhosphoUridine(H)@11 | 3510.815 | 3510.631 | 0.184073 | 3 |
| 23 | A0A0F6B7T9 | SEEPEVEEEYESDEDDTVDEER | PhosphoUridine(Y)@10 | 3268.43 | 3268.241 | 0.19096 | 3 |
| 24 | RL13 | AEYTPHVDTGDYIIVLNADK | PhosphoUridine(Y)@12 | 3148.697 | 3148.51 | 0.185498 | 3 |
| 25 | GAL1 | ALFAEIFGYPATHTIQAPGR | PhosphoUridine(Y)@9 | 2769.529 | 2769.347 | 0.181909 | 3 |
| 26 | YDIU | GILLGEQLLADGSTLDWHLK | PhosphoUridine(H)@18 | 3092.763 | 3092.604 | 0.159388 | 2 |

| | | | | | | | |
|---|---|---|---|---|---|---|---|
| 27 | EFTU | ELLSQYDFPGDDTPIVR | PhosphoUridine(Y)@6 | 2574.361 | 2574.183 | 0.177945 | 3 |
| 28 | A0A0F6AXS9 | ISALSDNGEHFSAK | PhosphoUridine(H)@10 | 2390.308 | 2390.125 | 0.182594 | 3 |
| 29 | SYI | GLSGFDSPYVPGWDCHGLPIELK | PhosphoUridine(Y)@9 | 3457.832 | 3457.652 | 0.180421 | 3 |
| 30 | PTA | DVLMEEIIANYHANTK | PhosphoUridine(Y)@11 | 2774.538 | 2774.345 | 0.193214 | 3 |
| 31 | FTSH | LAEEIIYGVEHVSTGASNDIK | PhosphoUridine(H)@11 | 3159.737 | 3159.547 | 0.189659 | 3 |
| 32 | SYM | GHEVNFICADDAHGTPIMLK | PhosphoUridine(H)@13 | 3138.659 | 3138.477 | 0.183771 | 3 |
| 33 | DEOB | IAAGMDGNADVIGAYAWAHELSSGK | PhosphoUridine(Y)@15 | 3417.8 | 3417.616 | 0.182658 | 3 |
| 34 | RL13 | AEYTPHVDTGDYIIVLNADK | PhosphoUridine(Y)@12 | 3148.706 | 3148.51 | 0.195082 | 3 |
| 35 | A0A0F6B033 | DGFVLGDGAGMLVLEEYEHAK | PhosphoUridine(Y)@17 | 3179.678 | 3179.498 | 0.180759 | 3 |
| 36 | SODM | SYTLPSLPYAYDALEPHFDK | PhosphoUridine(Y)@11 | 3240.746 | 3240.552 | 0.195406 | 3 |
| 37 | CH60 | GYLSPYFINKPETGAVELESPFILLADK | PhosphoUridine(Y)@6 | 4329.434 | 4329.263 | 0.172747 | 3 |
| 38 | OMPC | VDGLHYFSDDKGSDGDQTYMR | PhosphoUridine(Y)@6 | 3319.64 | 3319.459 | 0.180325 | 3 |
| 39 | ATPB | EGNDFYHEMTDSNVIDK | PhosphoUridine(H)@7 | 2927.462 | 2927.278 | 0.183543 | 3 |
| 40 | ARAA | VMSTGLQGGTSFMEDYTYHFEK | PhosphoUridine(Y)@18 | 3441.728 | 3441.54 | 0.187638 | 3 |
| 41 | 6PGD | AASDEYHWDLNYGEIAK | PhosphoUridine(H)@7 | 2895.511 | 2895.321 | 0.18977 | 3 |
| 42 | DEOB | FGDVGSDTLGHIAEACAK | PhosphoUridine(H)@11 | 2761.467 | 2761.288 | 0.178746 | 3 |
| 43 | RL10 | DTFVGPTLIAYSMEHPGAAAR | PhosphoUridine(Y)@11 | 2813.491 | 2813.304 | 0.187386 | 3 |
| 44 | A0A0F6B748 | SFPNICYFALSPEDEAR | PhosphoUridine(Y)@7 | 2625.326 | 2625.14 | 0.186466 | 3 |
| 45 | ARAB | ALAVDCATGDEIATSVEWYPR | PhosphoUridine(Y)@19 | 2933.493 | 2933.31 | 0.183679 | 3 |
| 46 | RRF | ASPSLLDGIVVEYYGTPTPLR | PhosphoUridine(Y)@14 | 2857.594 | 2857.409 | 0.184227 | 3 |
| 47 | RS11 | ITNITDVTPIPHNGCRPPK | PhosphoUridine(H)@12 | 3043.712 | 3043.541 | 0.169854 | 3 |
| 48 | RL13 | AEYTPHVDTGDYIIVLNADK | PhosphoUridine(Y)@12 | 3147.686 | 3147.526 | 0.160821 | 3 |
| 49 | A0A0F6AYJ0 | ILEVLQEPDNHHVSAEDLYK | PhosphoUridine(H)@12 | 3262.778 | 3262.601 | 0.17713 | 3 |
| 50 | FABD | SVEFIAAQGVEHLYEVGPGK | PhosphoUridine(Y)@14 | 3043.7 | 3043.515 | 0.184889 | 3 |
| 51 | FABH | VAVTELAHIVDETLAANNLDR | PhosphoUridine(H)@8 | 2873.58 | 2873.411 | 0.168654 | 3 |
| 52 | NFSB | GYTSLVVVPVGHHSVEDFNAGLPK | PhosphoUridine(H)@13 | 3435.92 | 3435.732 | 0.187892 | 3 |
| 53 | RPOC | MGHIELASPTAHIWFLK | PhosphoUridine(H)@12 | 2864.643 | 2864.455 | 0.187871 | 3 |
| 54 | YDIU | AREGSDYTRTFRMLSHTEQQSASSPLR | PhosphoUridine(Y)@7 | 3416.615 | 3416.521 | 0.093613 | 3 |
| 55 | CLPB | GYEIHISDEALKLLSANGYDPVYGARPLK | PhosphoUridine(Y)@23 | 3494.651 | 3494.676 | -0.02326 | 3 |
| 56 | ATPA | DRGEDALIIYDDLSK | PhosphoUridine(Y)@10 | 2027.783 | 2027.872 | -0.08899 | 2 |

UMPylated sites were highlighted by gray boxes.

**Table S2 UMPylated proteins identified in YdiU-expressing *Salmonella***

| No | Protein | Full Name | Function | Modification Type |
|---|---|---|---|---|
| 1 | YdiU | UPF0061 family | Uncharacterized | PhosphoUridine(Y) PhosphoUridine(H) |
| 2 | DnaK | Heat shock 70 kDa protein | Chaperone | PhosphoUridine(Y) |
| 3 | GroEL | Heat shock 60 kDa protein | Chaperone | PhosphoUridine(Y) |
| 4 | HtpG | Heat shock 90 kDa protein | Chaperone | PhosphoUridine(H) |
| 5 | ClpB | Heat shock protein F84.1 | Chaperone | PhosphoUridine(Y) |
| 6 | Lon | ATP-dependent serine protease | protease | PhosphoUridine(H) |
| 7 | FtsH | ATP-dependent zinc protease | protease | PhosphoUridine(H) |
| 8 | RplM | 50S ribosomal protein L13 | ribosome | PhosphoUridine(Y) |
| 9 | RplJ | 50S ribosomal protein L10 | ribosome | PhosphoUridine(Y) |
| 10 | RpsQ | 30S ribosomal protein S17 | ribosome | PhosphoUridine(H) |
| 11 | RpsK | 30S ribosomal protein S11 | ribosome | PhosphoUridine(H) |
| 12 | IleS | Isoleucine--tRNA ligase | tRNA ligase | PhosphoUridine(Y) |
| 13 | MetG | Methionine--tRNA ligase | tRNA ligase | PhosphoUridine(H) |
| 14 | Frr | Ribosome-recycling factor | translational termination | PhosphoUridine(Y) |
| 15 | LepA | Elongation factor 4 | Elongation factor | PhosphoUridine(Y) |
| 16 | TufA | Elongation factor Tu | Elongation factor | PhosphoUridine(Y) |
| 17 | AtpA | ATP synthase subunit alpha | ATP synthase | PhosphoUridine(Y) |
| 18 | AtpD | ATP synthase subunit beta | ATP synthase | PhosphoUridine(Y) PhosphoUridine(H) |
| 19 | AtpG | ATP synthase subunit gamma | ATP synthase | PhosphoUridine(Y) |
| 20 | PurA | Adenylosuccinate synthetase | AMP biosynthesis | PhosphoUridine(Y) |
| 21 | RpoB | RNA polymerase subunit beta | RNA polymerase | PhosphoUridine(Y) |
| 22 | RpoC | RNA polymerase subunit beta’ | RNA polymerase | PhosphoUridine(H) |
| 23 | GlpQ | Glycerophosphodiester phosphodiesterase | glycometabolism | PhosphoUridine(H) |
| 24 | GapA | Glyceraldehyde-3-phosphate dehydrogenase | glycometabolism | PhosphoUridine(H) |
| 25 | GlmS | Glutamine--fructose-6-phosphate aminotransferase | glycometabolism | PhosphoUridine(H) |
| 26 | YeaD | Putative glucose-6-phosphate 1-epimerase | glycometabolism | PhosphoUridine(Y) |
| 27 | GND | 6-phosphogluconate dehydrogenase | glycometabolism | PhosphoUridine(H) |
| 28 | AraA | L-arabinose isomerase | glycometabolism | PhosphoUridine(Y) |
| 29 | AraB | Ribulokinase | glycometabolism | PhosphoUridine(Y) PhosphoUridine(H) |
| 30 | AraD | L-ribulose-5-phosphate 4-epimerase | glycometabolism | PhosphoUridine(Y) |
| 31 | GalK | Galactokinase | glycometabolism | PhosphoUridine(Y) |
| 32 | FabF | 3-oxoacyl-[acyl-carrier-protein] synthase 2 | Lipid metabolism | PhosphoUridine(Y) |
| 33 | FabD | Malonyl CoA-acyl carrier protein transacylase | Lipid metabolism | PhosphoUridine(Y) |
| 34 | FabH | 3-oxoacyl-[acyl-carrier-protein] synthase 3 | Lipid metabolism | PhosphoUridine(H) |

| | | | | |
|---|---|---|---|---|
| 35 | PNP | Polyribonucleotide nucleotidyltransferase | mRNA degradation | PhosphoUridine(H) |
| 36 | MetK | S-adenosylmethionine synthase | AdoMet synthase | PhosphoUridine(H) |
| 37 | MurA | UDP-N-acetylglucosamine 1-carboxyvinyltransferase | Cell wall formation | PhosphoUridine(H) |
| 38 | pta | Phosphate acetyltransferase | acetyl-CoA biosynthesis | PhosphoUridine(Y) |
| 39 | DeoB | Phosphopentomutase | 5-phospho-alpha-D-ribose1-diphosphate biosynthesis | PhosphoUridine(Y) PhosphoUridine(H) |
| 40 | pflB | Pyruvate formate lyase I | pyruvate fermentation | PhosphoUridine(Y) |
| 41 | SodA | Superoxide dismutase, Sod_Fe | antioxidant | PhosphoUridine(Y) |
| 42 | SodB | Superoxide dismutase | antioxidant | PhosphoUridine(Y) |
| 43 | nfnB | Oxygen-insensitive NAD(P)H nitroreductase | antioxidant | PhosphoUridine(H) |
| 44 | Fur | Ferric uptake regulation protein | transcription factor | PhosphoUridine(H) |
| 45 | LpoA/ yraM | Penicillin-binding protein activator | enzyme regulator activity | PhosphoUridine(Y) |
| 46 | OmpC | Outer membrane porin protein C | porin protein | PhosphoUridine(Y) |

**Table S3 Hydrogen bonds identified in YdiU-UTP-Mn and YdiU-ATP-Mn models**

| Donor | Acceptor | AvgDist. (Å) | AvgAng. (°) | Frec. (%) |
|---|---|---|---|---|
| **YdiU-UTP-Mn** | | | | |
| Arg177-NH2-HH22 | UTP-O1G | 2.71 | 164.45 | 99.84 |
| Arg170-NH1-HH12 | UTP-O3G | 2.71 | 159.61 | 99.64 |
| Arg170-NH2-HH22 | UTP-O1G | 2.77 | 161.17 | 94.84 |
| Arg87-NE-HE | UTP-O2B | 2.79 | 155.47 | 92.32 |
| UTP-O3-HO3 | Tyr71-OH | 2.81 | 161.60 | 80.81 |
| UTP-O2-HO2 | Gly84-O | 2.74 | 145.97 | 69.54 |
| Arg177-NH1-HH12 | UTP-O2G | 2.83 | 155.28 | 61.82 |
| Gly120-N-H | UTP-O4A | 2.87 | 151.45 | 56.97 |
| Lys107-NZ-HZ3 | UTP-O3G | 2.72 | 161.23 | 42.55 |
| Gly86-N-H | UTP-O2 | 2.89 | 151.32 | 41.21 |
| Arg87-NH1-HH11 | UTP-O2B | 2.83 | 146.31 | 40.10 |
| **YdiU-ATP-Mn** | | | | |
| Arg170-NH2-HH22 | ATP-O3G | 2.79 | 160.11 | 92.95 |
| Arg177-NH2-HH22 | ATP-O3G | 2.78 | 152.95 | 76.21 |
| Arg170-NH1-HH12 | ATP-O1G | 2.87 | 162.74 | 61.22 |
| Arg87-N-H | ATP-O1B | 2.84 | 148.10 | 39.53 |
| Arg177-NH1-HH12 | ATP-O3G | 2.86 | 147.65 | 39.33 |
| Gly86-N-H | ATP-O5 | 2.90 | 150.85 | 26.49 |

**Table S4 NMPylated pepetides in WT and ΔYdiU with and without heat shock.**

### WT with heat treatment

| UMP | Pro | Sequence | Modifications | P. MW | T.MW | dM | z |
|---|---|---|---|---|---|---|---|
| 1 | OmpC | LAAQYSQTYNATRFGTSNGSNPSTSYGFANK | PhosphoUridine(Y)@9 | 3608.504 | 3608.548 | -0.04351 | 3 |
| 2 | OmpD | LDLYGKVHAQHYFSDDNGSDGDKTYAR | PhosphoUridine(H)@11 | 3378.482 | 3378.41 | 0.071137 | 3 |
| 3 | GyrB | IHRQIYEHGVPQAPLAVTGDTDK | PhosphoUridine(H)@8 | 2851.419 | 2851.318 | 0.101439 | 2 |
| 4 | TlpA | PATYEPEQIIEAGLALQAEGR | PhosphoUridine(Y)@4 | 2561.176 | 2561.169 | 0.007269 | 3 |
| 5 | Dps | YAVVANDVRKAIGEAKDEDTADIFTAASR | PhosphoUridine(Y)@1 | 3401.786 | 3401.578 | 0.209739 | 3 |
| 6 | STM14_4308 | GHTLQQLDSIISAKGQTAYSSIVLGK | PhosphoUridine(Y)@19 | 3020.642 | 3020.486 | 0.155442 | 3 |
| 7 | YdiU | MLSHTEQQSASSPLR | PhosphoUridine(H)@4 | 1992.866 | 1992.825 | 0.041308 | 2 |
| 8 | PflB | QYVTALNVIHYMHDKYSYEASLMALHDR | PhosphoUridine(H)@13 | 3674.477 | 3674.621 | -0.14341 | 3 |
| AMP | Pro | Sequence | Modifications | P. MW | T.MW | dM | z |
| 1 | FliC | FNSAITNLGNTVNNLTSAR | Phosphoadenosine(T)@6 | 2335.141 | 2335.071 | 0.070752 | 2 |
| 2 | ClpB | GTLTDLLKSAGATTANITQAIEQMR | Phosphoadenosine(K)@8 | 2948.507 | 2948.406 | 0.101054 | 3 |
| 3 | GapA | AGIALNDNFVKLVSWYDNETGYSNK | Phosphoadenosine(K)@11 | 3147.488 | 3147.397 | 0.090255 | 3 |
| 4 | OmpC | TTSKRTADQNNTANAR | Phosphoadenosine(K)@4 | 2076.964 | 2076.909 | 0.055173 | 3 |
| 5 | NuoG | AIAHALDNTAPAVDGIDSDLQNK | Phosphoadenosine(T)@9 | 2677.323 | 2677.213 | 0.109406 | 3 |
| 6 | Kgd | LNVLINVLGKKPQDLFDEFAGK | Phosphoadenosine(K)@10 | 2787.373 | 2787.4 | -0.0266 | 3 |
| 7 | OmpD | TRLAFAGLKFADYGSFDYGR | Phosphoadenosine(K)@9 | 2583.169 | 2583.17 | -0.00211 | 3 |
| 8 | TypA | RGKVKPNQQVTIIDSEGK | Phosphoadenosine(K)@3 | 2325.189 | 2325.159 | 0.029546 | 2 |
| 9 | STM14_2170 | LFAGAGARVVAIQDHTATLFNATGIDMK | Phosphoadenosine(H)@15 | 3216.539 | 3216.554 | -0.01451 | 3 |
| 10 | SspA | TLFSGPTDIYSHQVR | Phosphoadenosine(T)@1 | 2048.991 | 2048.911 | 0.08034 | 3 |
| GMP | Pro | Sequence | Modifications | P.MW | T.MW | dM | z |
| 1 | GroEL | EIELEDKFENMGAQMVK | Phosphoguanosine(K)@7 | 2355.023 | 2354.991 | 0.032641 | 2 |

| | | | | | | | |
|---|---|---|---|---|---|---|---|
| 2 | HtpG | EGPAEDHANQEAIAKLLRFASTHTDSSAQTVSLEDYVSR | Phosphoguanosine(K) @15 | 4604.135 | 4604.084 | 0.052664 | 3 |
| 3 | RpsA | NRAISLSVRAKDEADEKDAIATVNK | Phosphoguanosine(K) @11 | 3059.405 | 3059.467 | -0.06269 | 3 |
| 4 | AceF | VTAEQSLITVEGDKASMEVPAPFAGTVK | Phosphoguanosine(K) @14 | 3235.586 | 3235.511 | 0.075692 | 3 |
| 5 | GapA | GANFDKYEGQDIVSNASCTTNCLAPLAK | Phosphoguanosine(K) @28 | 3404.615 | 3404.444 | 0.172046 | 3 |
| 6 | OmpA | GIPSDKISARGMGESNPVTGNTCDNVKPR | Phosphoguanosine(K) @6 | 3417.587 | 3417.519 | 0.067544 | 3 |
| 7 | OmpA | LISKGIPSDKISARGMGESNPVTGNTCDNVKPR | Phosphoguanosine(K) @10 | 3842.756 | 3842.819 | -0.06218 | 3 |
| 8 | AdhE | LAHVDKLSEDAFDDQCTGANPR | Phosphoguanosine(K) @6 | 2803.174 | 2803.166 | 0.008015 | 3 |
| **ΔYdiU with heat treatment** | | | | | | | |
| AMP | Pro | Sequence | Modifications | P.MW | T.MW | dM | z |
| 1 | TufA | PHVNVGTIGHVDHGK | Phosphoadenosine(T) @7 | 1894.871 | 1894.859 | 0.012125 | 3 |
| GMP | Pro | Sequence | Modifications | P.MW | T.MW | dM | z |
| 1 | TktA | IGFGSPNKQGTHDSHGAPLGDAEIALTR | Phosphoguanosine(H) @12 | 3190.406 | 3190.458 | -0.05315 | 3 |
| 2 | AceF | VGDKVEAEQSLITVEGDKASMEVPSPQAGVVK | Phosphoguanosine(K) @18 | 3657.795 | 3657.723 | 0.071806 | 4 |
| 3 | ArnA | ANAAEIYNKDGNKLDLFGKVDGLHYFSDDKGSDGDQTYMR | Phosphoguanosine(K) @19 | 4812.038 | 4812.119 | -0.0806 | 3 |
| 4 | PtsI | QVIDASHAEGKWTGMCGELAGDER | Phosphoguanosine(K) @11 | 2961.215 | 2961.217 | -0.00177 | 3 |
| 5 | TufA | AIDKPFLLPIEDVFSISGR | Phosphoguanosine(K) @4 | 2461.289 | 2461.204 | 0.085004 | 3 |
| **WT without heat treatment** | | | | | | | |
| GMP | Pro | Sequence | Modifications | P.MW | T.MW | dM | z |
| 1 | TufA | AIDKPFLLPIEDVFSISGR | Phosphoguanosine(K) @4 | 2461.289 | 2461.204 | 0.085004 | 3 |
| 2 | ArnA | HPLRCHFPPFAGFQVVESR | Phosphoguanosine(H) @6 | 2641.222 | 2641.18 | 0.04183 | 3 |

UMPylated sites were highlighted by gray boxes. P. MW: Prec MW; T.MW:Theor MW; dM: dMass; Z: Theor z.

**Table S5. Strains and plasmids used in this study**

| Strains | Relevant characteristic(s) | Source or reference |
|---|---|---|
| **Wild type *Salmonella*** | **ATCC14028 (No resistance)** | **Laboratory stock** |
| **Wild type *Salmonella* (Amp$^+$)** | **with pBad24 control (Amp$^+$)** | **This study** |
| **ΔYdiU** | **ydiU knockout strain (No resistance)** | **This study** |
| **ΔYdiU (Amp$^+$)** | **ΔYdiU with pBad24 control (Amp$^+$)** | **This study** |
| **pYdiU** | **ΔYdiU with YdiU$^{475}$/ pBAD24 (Amp$^+$)** | **This study** |
| **pYdiU D256A** | **ΔYdiU with YdiU$^{475}$ D256A/ pBAD24 (Amp$^+$)** | **This study** |
| ***E. coli* BL21(DE3)** | **T7 expression host** | **Laboratory stock** |
| ***E. coli* BL21(DE3) ΔYdiU** | **YdiU knockout strain of *E. coli* BL21(DE3)** | **This study** |
| ***E. coli* K12** | **Substr.MG1655, genome template** | **Laboratory stock** |
| **Plasmids** | **Relevant characteristic(s)** | **Source or reference** |
| **pBad24** | **Vector pBad24 control Amp$^+$** | **Laboratory stock** |
| **YdiU$^{475}$/pBad24** | ***Salmonella* YdiU$^{1\text{-}475aa}$ cloned into pBad24** | **This study** |
| **YdiU$^{475}$ D256A/pBad24** | ***Salmonella* YdiU$^{1\text{-}475aa}$ D256A cloned into pBad24** | **This study** |
| **pGL01** | **Expression Vector Amp$^+$** | **Laboratory stock** |
| **YdiU$^{fl}$/pGL01** | ***E.coli* YdiU full-length cloned into pGL01** | **This study** |
| **YdiU$^{475}$/pGL01** | ***E.coli* YdiU 1-475aa cloned into pGL01** | **This study** |
| **YdiU$^{477}$/pGL01** | ***E.coli* YdiU 1-477aa cloned into pGL01** | **This study** |
| **YdiU$^{478A}$/pGL01** | ***E.coli* YdiU full-length adding an Alanine in C-terminal, cloned into pGL01** | **This study** |
| **YdiU$^{478AA}$/pGL01** | ***E.coli* YdiU full-length adding two Alanines in C-terminal, cloned into pGL01** | **This study** |
| **GroEL191-376aa/pGL01** | ***E.coli* GroEL191-376aa cloned into pGL01** | **This study** |
| **DnaK /pGL01** | ***E.coli* DnaK 1-636aa cloned into pGL01** | **This study** |
| **HtpG/pGL01** | ***E.coli* HtpG 1-624aa cloned into pGL01** | **This study** |

| | | |
|---|---|---|
| **ClpB/pGL01** | ***E.coli* ClpB 1-857aa cloned into pGL01** | **This study** |
| **GrpE/pGL01** | ***E.coli* GrpE 38-197aa cloned into pGL01** | **This study** |
| **GrxA/pGL01** | ***E.coli* GrxA 1-85aa cloned into pGL01** | **This study** |
| **GroEL Y199F/pGL01** | ***E.coli* GroEL 191-376aa Y199F cloned into pGL01** | **This study** |
| **GroEL Y203F/pGL01** | ***E.coli* GroEL 191-376aa Y203F cloned into pGL01** | **This study** |
| **GroEL Y199F-Y203F/pGL01** | ***E.coli* GroEL 191-376aa Y199F-Y203F cloned into pGL01** | **This study** |
| **GroEL Y199F-Y203F-Y360F/pGL01** | ***E.coli* GroEL 191-376aa Y199F-Y203F-Y360F cloned into pGL01** | **This study** |
| **YdiU Y71A/pGL01** | ***E.coli* YdiU Y71A cloned into pGL01** | **This study** |
| **YdiU K107A/pGL01** | ***E.coli* YdiU K107A cloned into pGL01** | **This study** |
| **YdiU D119A/pGL01** | ***E.coli* YdiU D119A cloned into pGL01** | **This study** |
| **YdiU E130A/pGL01** | ***E.coli* YdiU E130A cloned into pGL01** | **This study** |
| **YdiU N244A/pGL01** | ***E.coli* YdiU N244A cloned into pGL01** | **This study** |
| **YdiU D246A/pGL01** | ***E.coli* YdiU D246A cloned into pGL01** | **This study** |
| **YdiU D246N/pGL01** | ***E.coli* YdiU D246N cloned into pGL01** | **This study** |
| **YdiU N247A/pGL01** | ***E.coli* YdiU N247A cloned into pGL01** | **This study** |
| **YdiU D256A/pGL01** | ***E.coli* YdiU D256A cloned into pGL01** | **This study** |

### Table S6. Data collection and refinement statistics

| | Se-Met-YdiU-AMPPNP | YdiU-AMP/imidodi phosphonic acid | Apo-YdiU | YdiU-AMPNPP with $Mn^{2+}$ |
|---|---|---|---|---|
| Protein Data Bank ID | 6INY | 6III | 6K20 | 6LNA |
| **Data collection** | | | | |
| Space group | $P6_122$ | $P6_122$ | $P2\ 2_1\ 2_1$ | *I121* |
| Cell dimensions | | | | |
| *a, b, c* (Å) | 71.64, 71.64, 364.36 | 71.42, 71.42, 363.45 | 42.53, 117.35, 147.79 | 112.43,73.54,122.65 |
| α, β, γ (°) | 90.00, 90.00, 120.00 | 90.00, 90.00, 120.00 | 90.00, 90.00, 90.00 | 90.00, 102.49, 90.00 |
| Wavelength (Å) | 0.9791 | 0.9791 | 0.9791 | 0.9791 |
| Resolution (Å) | 50.00-2.00 (2.07-2.00)* | 50.00-2.11 (2.19-2.11) | 27.89-2.58（2.69-2.58） | 27.44-1.70 (1.73-1.70) |
| ⟨ $I/\sigma(I)$⟩ | 71.83 (3.37) | 46.77 (5.27) | 20.5 (9.1) | 12.3 (5.3) |
| Completeness (%) | 99.5 (98.9) | 99.8 (99.9) | 93.6 (78.0) | 91.5 (96.4) |
| Redundancy | 27.6 (28.1) | 15.1 (16.3) | 11 (10.7) | 15.1 (16.3) |
| $CC_{1/2}$ | 0.992 (0.836) | 0.929 (0.996) | 0.997 (0.996) | 0.991 (0.845) |
| $R_{pim}$ | 0.020 (0.252) | 0.039 (0.175) | 0.028 (0.084) | 0.053 (0.221) |
| **Refinement** | | | | |
| Resolution (Å) | 30.91-2.00 | 30.93-2.11 | 27.89-2.58 | 27.44-1.70 |
| No. reflections | 38863 | 32104 | 22642 | 97925 |
| $R_{work}$ / $R_{free}$ (%) | 20.55/24.45 | 20.55/25.30 | 18.4/21.9 | 15.81/18.48 |
| No. atoms | | | | |
| Protein | 3710 | 3701 | 3899 | 7463 |
| Ion | 2 $Mg^{2+}$ | 2 $Mg^{2+}$ | 1 $Mg^{2+}$ | 4 $Mn^{2+}$ 2$Ca^{2+}$ |
| Ligands | 31 | 32 | | 64 |
| Water | 118 | 120 | 161 | 1662 |
| *B*-factors | | | | |
| Protein | 64.51 | 60.70 | 38.71 | 10.27 |
| $Mg^{2+}$ | 47.78 | 60.59 | 47.95 | |
| $Mn^{2+}$ | | | | 6.5 |
| $Ca^{2+}$ | | | | 15 |
| Ligands | 55.44 | 55.58 | | 5.5 |
| Water | 57.71 | 54.17 | 40.40 | |
| RMS deviations | | | | |
| Bond lengths (Å) | 0.015 | 0.007 | 0.005 | 0.004 |
| Bond angles (°) | 1.401 | 0.890 | 0.666 | 0.753 |

*values in parentheses are for highest-resolution shell.
